# Using gene complementation to identify a SulP-family bicarbonate transporter in an N2-fixing cyanobacterial endosymbiont of an open ocean diatom

**DOI:** 10.1101/2024.01.10.573558

**Authors:** Mercedes Nieves-Morión, Rubén Romero-García, Sepehr Bardi, Luis López-Maury, Martin Hagemann, Enrique Flores, Rachel A. Foster

## Abstract

Diatom-Diazotrophic Associations (DDAs) contribute significantly to new and primary production in the world’s oceans, yet the understanding of how production is sustained is poorly resolved. These symbioses involve diatoms and N_2_-fixing, heterocyst-forming cyanobacteria of the genus *Richelia*, both partners being photosynthetic. *Richelia euintracellularis* resides in the cytoplasm of *Hemiaulus hauckii*, whereas *Richelia intracellularis* is periplasmic in *Rhizosolenia clevei*. In the ocean, bicarbonate is taken up by phytoplankton to provide CO_2_ for photosynthesis. The genomes of both *Richelia* endobionts (ReuHH01 and RintRC01, respectively) contain genes encoding SulP-family proteins, which are oxyanion transporters. To study the possible involvement of these transporters in bicarbonate uptake, we used complementation of a *Synechocystis* sp. PCC 6803 mutant with its five CO_2_ uptake systems inactivated, which is unable to grow in air levels of CO_2_. Three genes from RintRC01 and one gene and a DNA fragment containing four partial gene sequences from ReuHH01 were chemically synthesized, cloned under the control of a strong gene promoter and incorporated in the chromosome of the *Synechocystis* mutant. One gene from RintRC01, RintRC_3892, complemented the *Synechocystis* mutant to grow with air levels of CO_2_ or with low bicarbonate concentrations. The complemented strain showed strong sodium-dependent, low affinity bicarbonate uptake, which, together with phylogenetic analyses, identified RintRC_3892 as a BicA protein. Additionally, RintRC_3892 transcripts were consistently detected in environmental samples from three ocean basins. No evidence for a bicarbonate transporter was found, however, for ReuHH01, suggesting different strategies for inorganic carbon uptake in the periplasmic and cytoplasmic endobionts.

## INTRODUCTION

The oceans play a main role in the carbon (C) cycle by taking up a significant amount of atmospheric CO_2_ and represent a natural C buffer in the biosphere [1]. In global oceans the total mass of C consists mostly of dissolved inorganic carbon (DIC), about 60 times greater than its atmospheric mass, and pools of particulate and dissolved organic carbon (POC and DOC, respectively) [2-4]. DIC includes three aqueous species: CO_2_, bicarbonate (HCO_3_^-^) and carbonate (CO_3_^2-^), which differ in their proportion according to the pH of the environment. Estimation of total DIC shows an increase associated with rising inputs of anthropogenic CO_2_ resulting in both spatial and temporal variability across the global ocean [5]. Marine phytoplankton living in the euphotic zone (sunlight zone) of the oceans perform photosynthesis using DIC to produce organic matter. Because the majority of DIC in the ocean is bicarbonate, phytoplanktonic organisms that actively transport bicarbonate across their cell membranes present a competitive advantage when inorganic C concentrations are limiting. Indeed, many cyanobacterial and algal species can rapidly acclimate to changes in the total concentration of DIC by the induction of CO_2_-concentrating systems (CCMs) [6].

Diatoms are a group of eukaryotic phytoplankton that are responsible for about 20% of the primary production on Earth [7] and are important contributors to the biological C pump that sequesters C to the deep ocean. Diatoms take up both CO_2_ and bicarbonate and concentrate DIC in their cells as part of the CCM [8-10]. Different transporters can actively concentrate DIC from the surrounding environment into the cell, where DIC enters the chloroplast compartment and carbonic anhydrases convert bicarbonate into CO_2_ that is fixed by ribulose 1-5-bisphosphate carboxylase/oxygenase (RubisCO) within the pyrenoid [11]. Additionally, some diatom species are supposed to use the plant-type C_4_ pathway for DIC accumulation and photosynthetic C fixation at low CO_2_ [12].

Carbon and nitrogen (N) metabolism are closely coupled in diatoms [13, 14]. Nitrogen availability in marine ecosystems varies over several spatial and temporal scales [15, 16] regulating the biological C pump [17]. Most diatoms reside in coastal zones where nutrients are high, while only a few genera of diatoms that live in symbiosis with N_2_-fixing cyanobacteria can prosper in oligotrophic (nutrient-poor) areas of the oceans. These symbioses are known as Diatom Diazotrophic Associations (DDAs) and have an important role in the biosphere given their high rates of both CO_2_ and N_2_ fixation and high measured export of fixed C to the deep ocean [18]. Two important DDAs are the associations between the diatoms *Hemiaulus hauckii* and *Rhizosolenia clevei* with the heterocyst-forming cyanobacteria *Richelia euintracellularis* and *Richelia intracellularis*, respectively [19]. Interestingly, the cellular location and integration of the symbiont with its respective diatom host varies. For example, in *H. hauckii*, *R. euintracellularis* is a fully integrated endobiont, residing in the diatom cytoplasm [20], whereas the *R. intracellularis* is only a partially integrated endobiont as a periplasmic symbiont, i.e., located between the cytoplasmic membrane and the frustule of *Rh. clevei* [21]. Additionally, a third *Richelia, R. rhizosoleniae*, often co-occurs with the other *Richelia* spp. symbioses and is a facultative symbiont that attaches extracellularly to the diatom *Chaetoceros compressus* [19, 22].

Genome sequences are available for *R. euintracellularis* (strain ReuHH01), *R. intracellularis* (strain RintRC01) and *R. rhizosoleniae* (strain RrhiSC01) [23, 24], as well as for several environmental metagenome assembled genomes (MAGs) [19]. Similar to other cyanobacteria, all the *Richelia* symbionts possess genes encoding several CCM components such as β-carboxysomes, RubisCO, and carboxysomal carbonic anhydrases, as well as a CO_2_ trapping system homologous to the cyanobacterial NDH-1_3_ complex [25]. Therefore, they are predicted to use CO_2,_ which is converted into bicarbonate by the NDH-1_3_ complex and then finally fixed by RubisCO inside the carboxysome. Indeed, high rates of C fixation have been reported from field studies when the DDAs are present [26-28]. Interestingly, the genome of the external and facultative symbiont (RrhiSC01) contains genes encoding proteins showing similarity to low- and high-affinity bicarbonate transporters, whereas genomes of both endobionts lack genes for high affinity bicarbonate transporters identified in other cyanobacteria [25], such as the Na^+^-dependent SbtA protein and the ABC-type Cmp transporter [29]. However, the *Richelia* endobiont genomes contain homologues to the SulP-family transporters, which includes the low-affinity, Na^+^- dependent bicarbonate transporter BicA, first described in *Synechococcus* sp. PCC 7002 [30]. SulP-family transporters are generally involved in the uptake of anions (frequently oxyanions) such as sulfate, nitrate, molybdate and, relevant here for autotrophy, bicarbonate, (https://www.tcdb.org/search/result.php?tc=2.A.53). Thus, some of the SulP-family transporters present in the endobionts could be bicarbonate transporters, but sequence homology is not sufficient to resolve neither their function nor specific substrate.

Here, we addressed the possible bicarbonate transporters identified as SulP-family transporters from the two endobionts, *R. euintracellularis* (ReuHH01) and *R. intracellularis* (RintRC01), in order to investigate whether they could function in inorganic carbon exchange within their respective symbiosis. Because the DDAs have largely evaded long term cultivation [31], and the endosymbionts appear unable of independent growth [19], we have used heterologous gene expression to investigate the transporter function and substrate affinity. The recipient organism used for the *Richelia* genes is the Δ5 mutant of the cyanobacterial model strain *Synechocystis* sp. PCC 6803, in which the five inorganic C uptake systems are inactivated [32]. Our study identifies one bicarbonate transporter from the periplasmic endobiont (RintRC01) which is functional and distinguishes the retention of an inorganic C uptake strategy that seems to be non-functional in the cytoplasmic endobiont (ReuHH01). Additionally, we report the expression of the gene encoding the newly identified bicarbonate transporter in wild populations in multiple ocean basins. Our results are consistent with the hypothesis that carbon metabolism in these cyanobacteria has simplified as a result of adaption to an endosymbiotic lifestyle.

## MATERIALS AND METHODS

### Strains and growth condition

*Synechocystis* sp. strain PCC 6803 was grown in BG11 medium (BG11 medium contains 0.189 mM Na_2_CO_3_), modified to contain ferric citrate instead of ferric ammonium citrate [33], at 30 °C in the light (ca. 25 to 30 µmol photons m^-2^ s^-1^) in shaken (100 rpm) liquid cultures. The *Synechocystis* Δ5 mutant (Δ*ndhD3/ndhD4/cmpA/sbtA/bicA*) [32] was grown in BG11C-medium (BG11 medium supplemented with 50 mM NaHCO_3_), bubbled with 1% (v/v) CO_2_ in air under light (50 µmol photons m^-2^ s^-1^), and the following antibiotics at the indicated concentrations: kanamycin (Km), 10 µg ml^-1^; streptomycin dihydrochloride pentahydrate (Sp), 5 µg ml^-1^; chloramphenicol (Cm), 5 µg ml^-1^; hygromycin (Hyg), 2 µg ml^-1^; and gentamicin (Gent), 1 µg ml^-1^. Cultures were diluted every 5-6 days. To select for a possible complementing gene, erythromycin (Em) at 10 µg/ml was added to the BG11C medium supplemented with 10 mM NaHCO_3_ and the five antibiotics described above. For tests on solid medium, BG11 medium was solidified with 1% (wt/v) purified Difco Bacto agar. Chlorophyll *a* (Chl) content of the cultures was determined by methanol extraction [34]. *E. coli* strain DH5α, used for plasmid constructions, was grown in Luria broth (LB) medium, supplemented when appropriate with antibiotics at standard concentrations [35].

### Analysis of DNA and protein sequences *in silico*

The search for sequences and data analysis to prepare constructs with the different genes of *Richelia* spp. was carried out using the Integrated Microbial Genomes (IMG) page of the Joint Genome Institute (https://img.jgi.doe.gov). To align the sequences of proteins, Clustal W (https://www.ebi.ac.uk/Tools/msa/clustalo/) was used. Phylogenetic analysis of SulP-like proteins from *Richelia* spp. was performed in Phylogeny.fr (http://www.phylogeny.fr) with default values (see Supplementary Methods for the ID numbers of the genes/proteins used).

### Construction of plasmids containing the symbiont genes

The five symbiotic genes, RintHH_3990-60 (1706 bp, including RintHH_3990, RintHH_3980, RintHH_3970 and RintHH_3960 and intervening sequences) and RintHH_20770 (1470 bp) from *Richelia euintracellularis* HH01, and RintRC_3409 (1470 bp), RintRC_3892 (1704 bp) and RintRC_4851 (1611 bp) from *Richelia intracellularis* RC01, were cloned in plasmid pNRSD_P*_cpcB_*_Ery [36] containing sequences of *nrsD* as DNA fragments for recombination, the promoter of *cpcB* (encoding C-phycocyanin beta subunit) and an erythromycin resistance cassette. The symbiotic genes were synthetized with 100% sequence verification by Integrated DNA Technologies (IDT, Leuven, Belgium) and provided in a pUCIDT plasmid. RintHH_3990-60, RintHH_20770 and RintRC_3409 from the pUCIDT plasmids were digested with BamHI and cloned into pNRSD_P*_cpcB_*_Ery, producing pMN21, pMN22 and pMN24, respectively. RintRC_4851 from pUCIDT was digested with NdeI and XhoI and cloned into pNRSD_P*_cpcB_*_Ery, producing the pMN23. RintRC_3892 was PCR-amplified from a synthetic RintRC_3892 sequence using primers RintRC_3892-5 and RintRC_3892-6 with NdeI and SphI restriction enzymes and cloned into pNRSD_P*_cpcB_*_Ery producing the plasmid pMN10. To verify the constructs and select the correct gene orientation, which is the same orientation of the *cpcB* promoter, PCR was carried out using primer P*_cpcB560_*_F_EcoRI and specific reverse primers for each gene: RintHH_3990-1, RintHH_20770-1, RintRC_3409-1 and RintRC_4851-1, RintRC_3892-2 (Supp. Table S1).

### Transformation of *Synechocystis* Δ5 mutant

To attempt complementation with the symbiotic genes, the *Synechocystis* Δ5 mutant was transformed with the plasmids carrying the genes of interest (pMN21, pMN22, pMN24, pMN23 and pMN10, respectively). Optical density at 750 nm (OD_750nm_) of the *Synechocystis* Δ5 mutant was set up between 0.6 and 1. Mutant cells were washed and resuspended with BG11C medium (BG11 supplemented with 10 mM NaHCO_3_), and 200 μl cells were mixed with 20 μl of the plasmid (about 200 ng DNA) and incubated in darkness overnight at room temperature. After that, transformed cells were incubated with 1 ml of BG11C medium for 4 h at 30 °C under low light condition. Subsequently, the cells were placed into BG11C agar plates for 48 h at 30 °C under low light. Transformed cells were then selected with erythromycin (Em) at 10 μg/ml. Em-resistant clones were isolated and their genetic structure was verified by PCR as described above. Strains bearing the corresponding construct were named RintHH_3990-60, RintHH_20770, RintRC_3409, RintRC_4851 and RintRC_3892, respectively.

### Growth tests on solid medium

For tests of growth on solid medium, *Synechocystis* sp. PCC 6803 (wild-type strain [WT], used as control) and the *Synechocystis* Δ5 mutant transformed with constructs carrying RintHH_3990-60, RintHH_20770, RintRC_3409, RintRC_4851 or RintRC_3892 were grown in BG11C-medium (10 mM of NaHCO_3_) liquid cultures with the appropriate antibiotics (at final concentration as described above) and air levels of CO_2_ in a shaker. *Synechocystis* Δ5 mutant (used as negative control) was grown as indicated above (50 mM NaHCO_3_ and bubbling with 1% [v/v] CO_2_). These cultures were centrifuged and washed three times with BG11_0_ medium (without any source of nitrogen and without added bicarbonate). The pellets were resuspended in BG11 medium and dilutions containing 1, 0.5, 0.25, 0.125, and 0.0625 µg Chl ml^-1^ were prepared. A 10-µl portion of each dilution was spotted on plates with BG11-agar or BG11-agar supplemented with 100 mM of NaHCO_3_ without antibiotics. The plates were incubated at 30 °C in the light.

### Growth tests on liquid medium

The growth rate constant (µ = [ln2]/t_d_, where t_d_ is the doubling time) was calculated from the increase in the optical density at 750 nm (OD_750nm_) of bubbled and shaken liquid cultures. Cultures of *Synechocystis* sp. PCC 6803 (wild-type strain used as control), the *Synechocystis* Δ5 mutant (used as negative control), and the *Synechocystis* Δ5 mutant transformed with constructs carrying RintHH_3990-60, RintHH_20770, RintRC_3409, RintRC_4851 or RintRC_3892 were inoculated at 0.1 µg Chl ml^-1^. Growth rate constants were determined under different conditions based on the NaHCO_3_ concentration: (i) 0, 1 and 10 mM of NaHCO_3_ were tested in cultures bubbled with air levels of CO_2_ and grown at 30 °C in the light, 50 µmol photons m^-2^ s^-1^; (ii) 1 and 10 mM of NaHCO_3_ were tested in shaken cultures (100 rpm) with air levels of CO_2_ and grown at 30 °C in the light (ca. 25 to 30 µmol photons m^-2^ s^-1^). Cultures were supplemented with the appropriate antibiotics at the concentration indicated earlier.

### 14C-bicarbonate uptake assay

To test bicarbonate transport in the *Synechocystis* strains, a ^14^C-bicarbonate uptake assay was performed. Bicarbonate uptake was routinely determined in 1-min assays with the indicated concentration of [^14^C]NaHCO_3_ at 30 °C in the light (about 75 μmol m^-2^ s^-1^) in BG11 medium or in a buffer in air-levels of CO_2_ and supplemented with NaCl at the indicated concentration. Cells that had been grown as indicated earlier and incubated for 18 h in BG11 medium bubbled with air were used at about 10 µg Chl ml^-1^. Solutions of ^14^C-labeled bicarbonate for the uptake assays were prepared by mixing unlabeled HNaCO_3_ and [^14^C]NaHCO_3_ at 55 mCi /mmol from American Radiolabeled Chemicals, Inc. To study the effect of Na^+^ (provided as NaCl) on the uptake of 0.11 mM [^14^C]NaHCO_3_ in *Synechocystis* WT and the Δ5 mutant complemented with RintRC_3892, cells prepared as described above were resuspended in 20 mM TES-KOH buffer, pH 8. To investigate the effect of the concentration of NaHCO_3_ on the uptake [^14^C]NaHCO_3_ in *Synechocystis* WT, Δ5 mutant and Δ5 mutant complemented with RintRC_3892, cells prepared as described above were resuspended in 20 mM TES-KOH buffer, pH 8, and supplemented with 30 mM NaCl (Fig. 3B). To test the bicarbonate uptake at pH 7 and pH 6 by *Synechocystis* WT and the strains carrying the *Richelia* genes, cells prepared as described above were incubated with 1 mM [^14^C]NaHCO_3_ in BG11/2 medium with 25 mM TES-KOH (pH 7) or MES-KOH (pH 6) buffer with a final Na^+^ concentration of about 18.5 mM. In every case, the assay was started by mixing cells and substrate. After 1 min (or different times in the experiment shown in Fig. 3A), the assay was finalized by a 25- to 50-fold dilution in ice-cold medium or buffer; the cells were harvested by filtration using 0.45-µm pore size filters, and the filters were washed three times with 2 to 3 mL of ice-cold medium or buffer. The radioactivity in the filters carrying the cells was then determined by scintillation counting; boiled cells were used as a blank.

### Environmental sampling and gene expression

Thirty-one whole water field samples (2-2.5 l) were collected for bulk RNA extraction from several locations in the North Atlantic and South China Sea (Supp. Table S3) using standard field methods (further details in Supplementary Methods). Total RNA was extracted using Qiagen RNAeasy Mini kit (Qiagen, Germany) following the manufacturer’s protocol with a few modifications described previously [22]. All samples were subjected to a 1-hour DNAse treatment (Qiagen RNase-free DNase kit), and final elution volume was 20 μl. Two to 5 μl of total RNA or diluted RNA (5 ng µl^-1^) was reverse transcribed (RT) using a commercially available kit (Superscript First Strand Synthesis System) and the cDNA (0.5-2 μl) was used as template in newly designed TaqMan quantitative polymerase chain reaction (qPCR) assays designed to quantify the expression of the genes encoding SulP-like proteins in *R. intracellularis.* The new assays were based on two *bicA* candidates: RintRC_3892 and RintRC_4851, and expression was normalized to *secA* (RintRC_6009). The target-specific TaqMAN (Applied Biosystems) oligonucleotides were designed and synthesized by IDT (Supp. Table S2; Suppl. Methods). Each probe was 5’labelled with a fluorescent reporter FAM (6-carboxyfluoreceom) and 3’ labelled with TAMRA (6-carboxytetramethulrhodamine) as a quenching dye. All qPCRs contained 12.5 μl of 2X TaqMAN buffer (Applied Biosystems), 10 μl of molecular grade water, 1.0 μl each of forward and reverse primers (10 μM), 0.5 ml probe (10 μM), and 1-2.0 μl cDNA template or separately 2 μl of water for no template controls and 2 μl of standards. Expression was quantified using duplicate standard curves of synthesized gBlocks (IDT) for each target made in an 8-point dilution series (10^8^ to 10^0^ gene copies per reaction) and run in parallel with all samples. Samples were run in triplicate reactions for cDNA and gene copies calculated based on the average cycle threshold (Ct) value and the standard curve for the appropriate assay. Samples that had 1 or 2 of the 3 replicates detected are reported as detected, not quantifiable (dnq); the limit of detection for the qPCR assays is 1-10 copies. Further details on the design and BLASTn analyses for testing the specificity of all oligonucleotides used in this study are described in full detail in the Supplementary Methods and results of in silico cross-reactivity tests are shown in Supp. Figure S7.

Gene expression profiles of the three SulP-like proteins from *R. intracellularis* RC01 (RintRC_3892, RintRC_4851, and RintRC_3904), were identified in a publicly available meta transcriptome dataset from the North Pacific Subtropical Gyre (NPSG) [37]. The genes were identified by using their original feature IDs as search queries in the Bacterial and Viral Bioinformatics Resource Center [38] and expression profiles for the SulP-like proteins were plotted in R [39].

## RESULTS

### SulP-like proteins are encoded in *Richelia* spp. genomes

The three species of *Richelia* whose genomes have been sequenced, *R. euintracellularis* HH01, *R. intracellularis* RC01, and *R. rhizosoleniae* SC01 (hereafter ReuHH01, RintRC01, and RrhiSC01, respectively), contain genes encoding SulP-like proteins [25]. Using the SulP protein BicA from *Synechococcus* sp. PCC 7002 and the SulP-like protein Alr1633 from *Anabaena* sp. PCC 7120 in BlastP analyses, four genes encoding full length SulP-like proteins were retrieved from RrhiSC01, three complete genes from RintRC01 (RintRC_3982, RintRC_4851, RintRC_3409), and one complete gene (RintHH_20770) as well as one possible gene split into four fragments (RintHH_3990-3980-3970-3960) from ReuHH01 (Supp. Fig. S1). To gain insight into the possible function of the *Richelia* SulP-like proteins, a phylogenetic analysis was carried out that included a representative number of non-metazoan SulP proteins from the Transporter Classification Database (TCDB) family 2.A.53 (https://tcdb.org/search/index.php?query=sulp&type=family). The sulfate and molybdate transporters, as well as one unusual nitrate transporter from cyanobacteria [40], segregated in the phylogenetic analysis from a group containing most of the SulP-like proteins encoded in the *Richelia* spp. genomes (Fig. 1). This group also contains BicA and SulP proteins associated to carbonic anhydrases (CAs), likely involved in bicarbonate uptake [41]. In this work, we were interested in identifying possible bicarbonate transporters of the endosymbiotic *Richelia*, i.e., RintRC01 (periplasmic) and ReuHH01 (cytoplasmic). We therefore focused on the three complete genes/proteins from RintRC01 and the two genes/proteins, one complete and another one split into four fragments, from ReuHH01 (Supp. Fig. S1). All were considered homologues to the SulP group that contains bicarbonate transporters (Fig. 1).

**Fig. 1.**
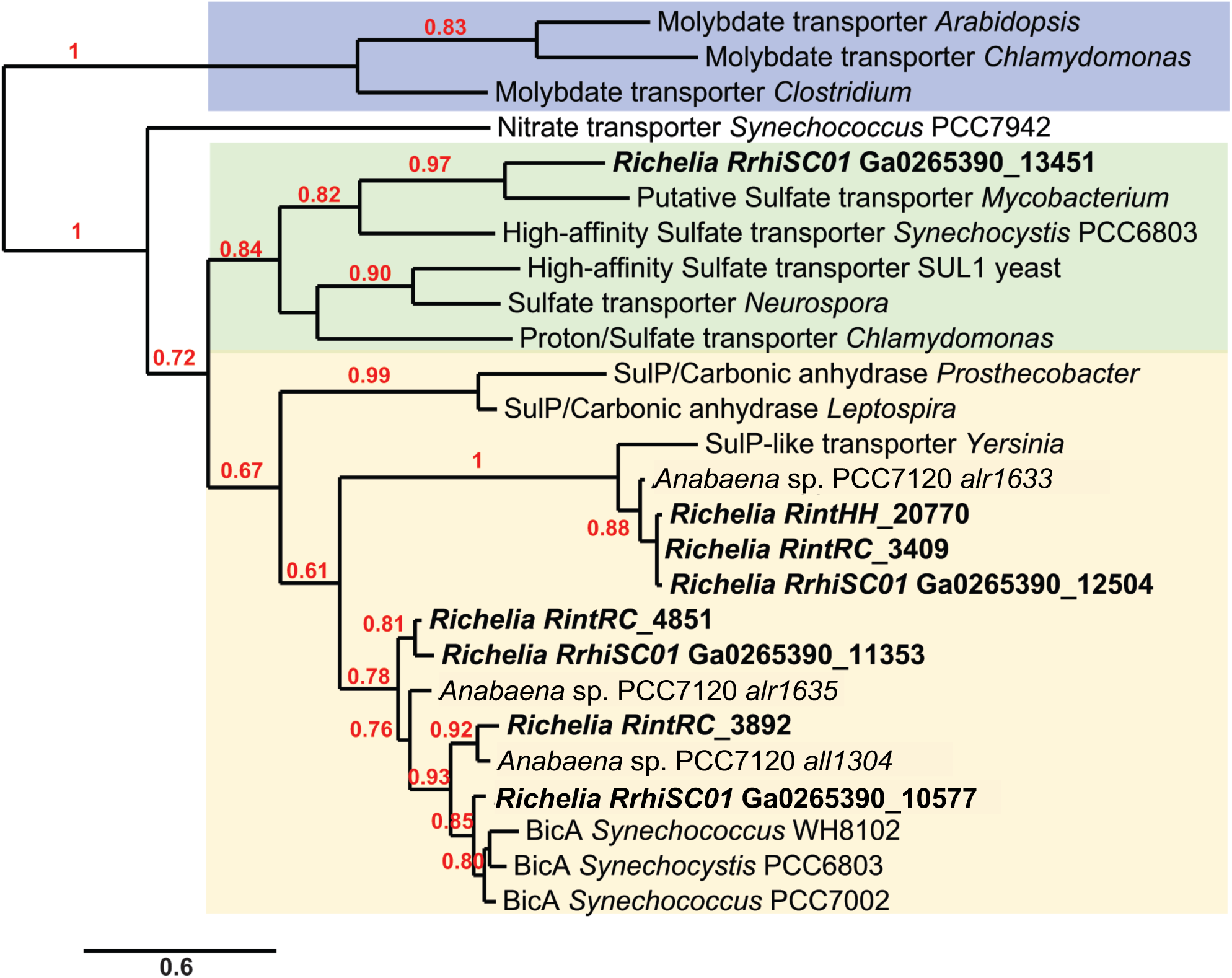
Phylogenetic analysis of SulP-like proteins from *Richelia* spp. Analysis performed in http://www.phylogeny.fr with default values. Number of bootstraps, 100. Scale bar is substitutions per site. See Supplementary Methods for ID gene/protein identification.

### *Synechocystis* Δ5 mutant is complemented by one *R. intracellularis* protein

To explore the possible role of the *Richelia* proteins in bicarbonate uptake, we tried complementation with each of the five *Richelia* genes in a mutant of *Synechocystis* sp. PCC 6803 that lacks all five CO_2_ and bicarbonate transporters [32]. This mutant, strain Δ5, is characterized by a high CO_2_-requiring (HCR) phenotype, because it is unable to grow in ambient air levels of CO_2_ [32, 42]. Using the available genomic information, *Richelia* genes were chemically synthesized and transferred to plasmid pNRSD_P*_cpcB_*_Ery, in which they were placed under control of the strong promoter P*_cpcB560_* upstream of an erythromycin-resistance gene (Fig. 2A; see also Supp. Fig. S2). In addition to cloning the following full gene candidates from *Richelia* RintRC01: RintRC_3892, RintRC_4851, RintRC_3409, we cloned the following gene candidates from *Richelia* ReuHH01 (formerly RintHH): RintHH_20770, and the whole genomic fragment covering the following annotated genes: RintHH_3990, RintHH_3980, RintHH_3970 and RintHH_3960, in an attempt to produce different polypeptides that might reconstitute a functional protein (Supp. Fig. S3). Hereafter, we refer to this latter sequence as RintHH_3990-60. The plasmids carrying the *Richelia* DNA sequences were transferred into *Synechocystis* strain Δ5 by genetic transformation and became incorporated into a neutral site (*nrsD*) of the chromosome by double homologous recombination. The *nrsD* gene determines nickel resistance and is non-essential under standard laboratory growth conditions (see Material and Methods) [43]. Transformants were selected on erythromycin-supplemented plates and further grown on plates of BG11 medium supplemented with 10 mM NaHCO_3_ and 10 µg Em ml^-1^. The presence of the P*_cpcB560_*-*sulP*-like gene construct in individual clones derived from the transformations was confirmed by PCR analysis (Supp. Fig. S4).

**Fig. 2.**
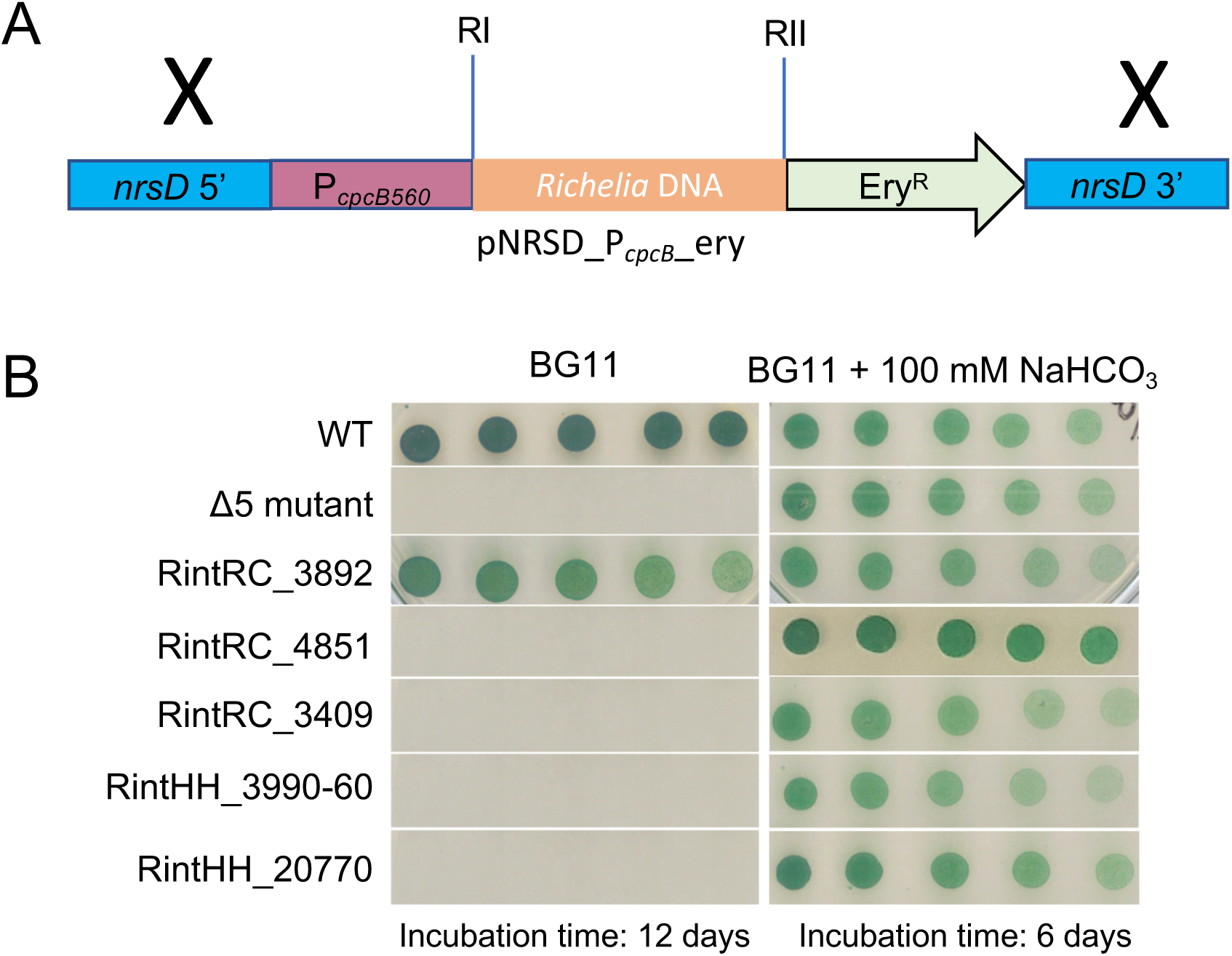
Complementation of *Synechocystis* strain D5 with *Richelia* spp. DNA (RintRC01 and ReuHH01). **(A)** Scheme of the construct used to introduce *Richelia* spp. DNA into the Δ5 mutant strain. The cloned DNA is located downstream from the P*_cpcB560_* promoter and adjacent to erythromycin-resistance (Ery^R^) gene, and the whole construct is inserted by double recombination in the non-essential *nrsD* gene. The X symbols designate approximate sites of recombination with the *Synechocystis* chromosome. Restriction sites: RI and RII are shown in detail in Fig. S3. **(B)** Photoautotrophic growth of wild-type *Synechocystis* (WT), the Δ5 mutant and derived strains carrying the indicated *Richelia* genes in solid BG11 medium (air levels of CO_2_; BG11 medium contains 0.189 mM Na_2_CO_3_) supplemented or not supplemented with 100 mM NaHCO_3_ and incubated as indicated (12 or 6 days).

The *Synechocystis* Δ5 mutant was routinely grown in liquid cultures of BG11 medium under high inorganic carbon conditions by the supplementation with 50 mM NaHCO_3_ and bubbled with 1% CO_2_-enriched air or on plates supplemented with 100 mM NaHCO_3_. The growth on plates of the *Synechocystis* Δ5 mutant transformed with the different *Richelia* constructs was first tested on solid BG11 medium using as controls the *Synechocystis* WT (hereafter WT) and Δ5 mutant strains (Fig. 2B). With air levels of CO_2_, only the WT and the Δ5 mutant transformed with RintRC_3892 could grow, whereas the Δ5 mutant and all the transformants with other *Richelia* RintRC01 genes (RintRC_4851, RintRC_3409) and the *Richelia* ReuHH01 genes (RintHH_20770, RintHH_3990-60) still showed the HCR phenotype and failed to grow (Fig. 2B). In contrast, all the strains could grow when the medium was supplemented with 100 mM NaHCO_3_ (Fig. 2B). Growth of the different strains was also tested in air-bubbled BG11 medium not supplemented or supplemented with 1 or 10 mM NaHCO_3_, and the growth rate constant was determined for the different strains (Table 1; Supp. Fig. S5). Without added bicarbonate, growth was observed for the WT and the Δ5 mutant carrying RintRC_3892 (0.51 ± 0.002 day^-1^ and 0.88 ± 0.006 day^-1^, respectively) (Table 1). With 1 mM NaHCO_3_, good growth was observed for the WT and the Δ5 mutant carrying RintRC_3892 (1.14 ± 0.2 day^-1^ and 1.07 ± 0.06 day^-1^, respectively). In both conditions, growth rate constants were significantly higher than for the Δ5 mutant (*P* < 0.001 by Student’s t test) (Table 1). In contrast, growth was observed for all strains, including the Δ5 mutant, when supplemented with 10 mM NaHCO_3_. Growth was also tested in shaken cultures of BG11 medium (ambient air levels of CO_2_); only the WT and the Δ5 mutant carrying RintRC_3892 grew in the presence of 1 mM NaHCO_3_, whereas all the strains could grow with 10 mM NaHCO_3_ (Supp. Fig S5). Combined the results show that RintRC_3892 encodes a protein that provides the *Synechocystis* Δ5 mutant with the capability to grow well in liquid or solid medium with ambient air levels of CO_2_ (which at chemical equilibrium should build a low concentration of bicarbonate) or with a low concentration (1 mM) of bicarbonate; i.e., the RintRC_3892 gene complements the HCR phenotype of the Δ5 mutant.

**Table 1.**
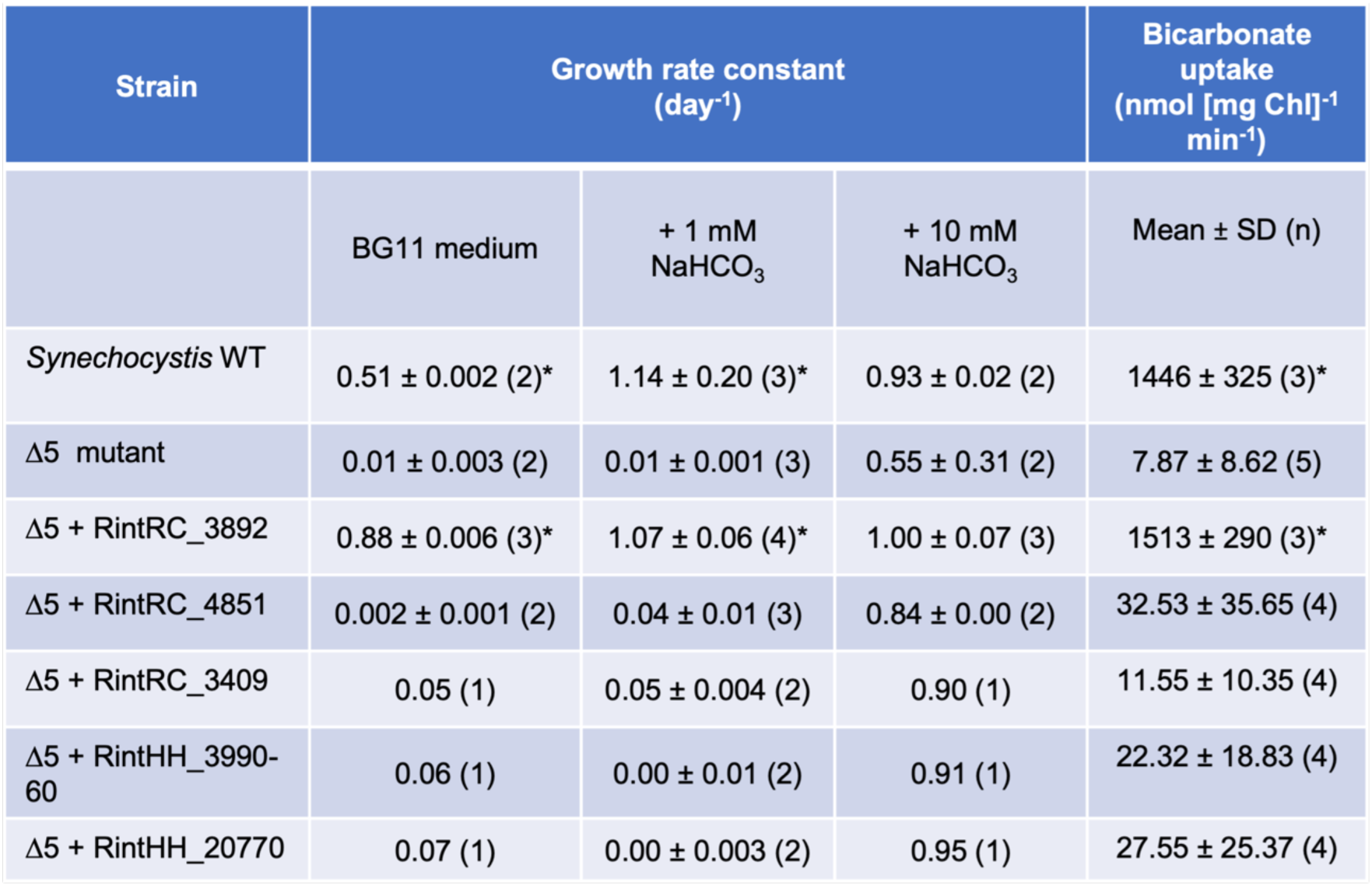
Growth test in bubbled liquid cultures and [^14^C]bicarbonate uptake in WT *Synechocystis* and 1′5 mutant transformed with *Richelia* genes. Growth was tested in autotrophic cultures with air-levels of CO_2_ and the indicated bicarbonate concentrations and described in Materials and Methods. The growth rate constant is presented for strains that could grow under the indicated conditions and was determined for the number of independent cultures shown in parenthesis. Uptake of 1.1 mM H^14^CO_3_ was tested in 1 min-assays performed at 30 °C in the light in BG11 medium supplemented with NaCl (final Na^+^ concentration ca. 31 mM), pH 8. The assays were carried out and the cells were prepared as described in Materials and Methods. Asterisks (*) denote significant differences in bicarbonate transport compared to the 1′5 mutant (*P* < 0.001 by Student’s t test).

### *Synechocystis* Δ5 transformed with SulP-like protein RintRC_3892 transports bicarbonate

To test directly transport of bicarbonate, we set up a bicarbonate uptake assay in which cells that had been incubated for 18 h in air-levels of CO_2_ were incubated in BG11 medium or a buffer and supplemented with ^14^C-labeled bicarbonate at 30 °C in the light (see Materials and Methods for details). As observed in the WT, bicarbonate supplied at 1 mM to cells in BG11 medium (pH 8) was efficiently taken up, and the rate of uptake progressively increased during a 2.5-min assay (Fig. 3A); this increase may reflect the time required for homogenization of the solution after mixing cells and substrate and subsequent transport of bicarbonate and trapping of CO_2_. The Δ5 mutant did not show any significant bicarbonate uptake, whereas the Δ5 mutant complemented with RintRC_3892 showed significant activity (*P* < 0.001 by Student’s t test) (Fig. 3A; Table 1).

**Fig. 3.**
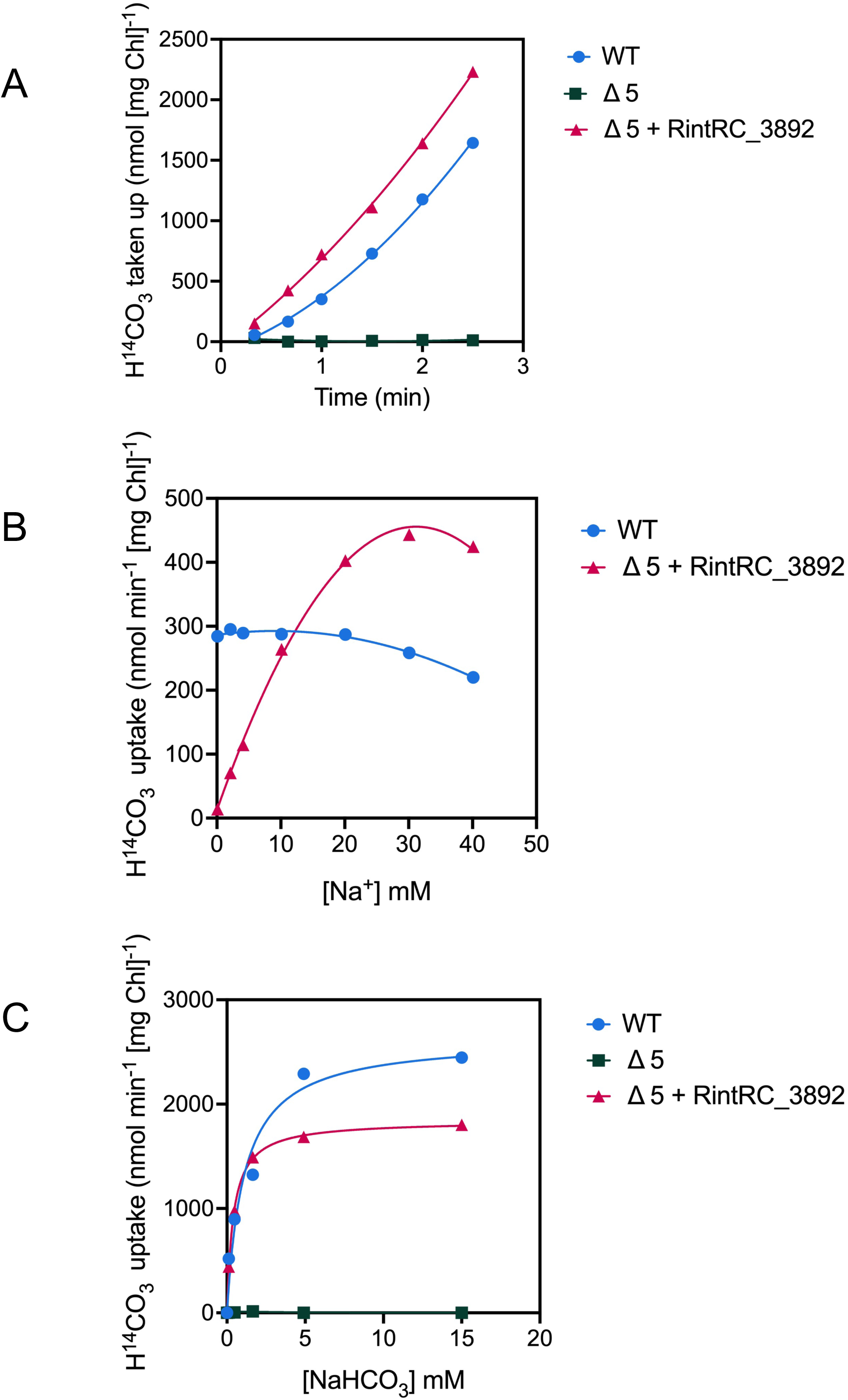
Uptake of ^14^C-bicarbonate by *Synechocystis* WT, the Δ5 mutant, and the Δ5 mutant transformed with RintRC_3892. **(A)** Uptake was tested with 1 mM bicarbonate (NaH^14^CO_3_) in BG11 medium supplemented with NaCl (final Na^+^ concentration, 31 mM) in cells that had been incubated for 18 h in BG11 medium (with antibiotics for the mutants) with air-levels of CO_2_. *Synechocystis* WT, blue circles; the Δ5 mutant, dark green squares; Δ5 mutant transformed with RintRC_3892, red triangles. **(B)** Na^+^-dependency of [^14^C]bicarbonate uptake in WT *Synechocystis* and mutant Δ5 complemented with RintRC_3892. Effect of Na^+^ (provided as NaCl) on the uptake of 0.11 mM NaH^14^CO_3_ in cells suspended in 20 mM TES-KOH buffer, pH 8. Shown is a representative experiment of assays with two independent cultures. **(C)** Bicarbonate-dependency of [^14^C]bicarbonate uptake in WT *Synechocystis* and mutant Δ5 complemented with RintRC_3892. Effect of the concentration of NaHCO_3_ on the uptake NaH^14^CO_3_ in cells suspended in 20 mM TES-KOH buffer, pH 8, supplemented with 30 mM NaCl (final sodium concentration increases above this value with added NaHCO_3_). Shown is a representative experiment of assays with three independent cultures. The assays were carried out and the cells were prepared as described in Materials and Methods.

To focus on initial transport rates, uptake of ^14^C-labeled bicarbonate was tested in 1-min assays with the different transformed strains. As shown earlier, the WT showed high ^14^C-bicarbonate uptake whereas the Δ5 mutant showed negligible uptake (Table 1). Among the Δ5 strains carrying the *Richelia* genes, the Δ5 mutant carrying RintRC_3892 showed high uptake of ^14^C-bicarbonate similar to WT rates, whereas the other transformants showed only slightly higher but not statistically significant increased activity than the Δ5 mutant (Table 1). In order to test whether a more acidic pH can stimulate the bicarbonate uptake activity, we ran assays at pH 6 and pH 7 resulting in similar results (Supp. Fig. S6). In conclusion, ^14^C-bicarbonate uptake assays are consistent with the liquid growth tests in which the medium was supplemented with 1 mM NaHCO_3_, showing that RintRC_3892 is a fully functional bicarbonate transporter.

Because the characterized SulP-family bicarbonate transporter, BicA, is sodium (Na^+^) dependent [30], we determined ^14^C-bicarbonate uptake in the RintRC_3892-complemented Δ5 mutant at a range of Na^+^ concentrations from 0.1 mM to 40 mM. The ^14^C-bicarbonate transport activity showed a strong sodium dependence (Fig. 3B), suggesting that RintRC_3892 mediates a Na^+^:HCO_3_^-^ co-transport as expected for a potential BicA homolog. The Δ5 mutant did not show ^14^C-bicarbonate uptake when tested at different Na^+^ concentrations (not shown), whereas the WT showed no Na^+^-dependency (Fig. 3B) or only a limited positive effect Na^+^ (not shown). It is likely that a significant part of ^14^C-bicarbonate uptake in the WT is mediated by Na^+^-independent transporters such as the ABC transporter Cmp.

We next tested the effect of ^14^C-bicarbonate concentration on uptake in the presence of Na^+^ (> 30 mM final concentration). The Δ5 mutant did not show uptake at any concentration tested, whereas the WT showed saturation kinetics likely reflecting the cumulative activity of its several bicarbonate transporters (Fig. 3C). The Δ5 mutant complemented with RintRC_3892 showed saturation kinetics with a *K*_0.5_ value of about 626 ± 244 µM (n = 3 independent cultures). Because RintRC_3892 is the only bicarbonate transporter in this strain, the determined *K*_0.5_ value is likely a fair indication of its affinity for bicarbonate.

As shown in Table 1 and Fig. 3, we frequently observed higher rates of ^14^C-bicarbonate uptake in the RintRC_3892-complemented Δ5 mutant than in the WT. This likely resulted from our strategy to use a very strong promoter (P*_cpcB560_*), which we chose to ensure expression in *Synechocystis* of the cloned *Richelia* genes.

### *Richelia* RintRC01 SulP-like transcripts are detected in environmental samples

In order to estimate the *in-situ* expression of some SulP-like proteins of *Richelia* RintRC01, we developed highly specific RT-qPCR assays to quantify gene transcripts of RintRC_3892 and RintRC_4851 in environmental samples (Supp. Fig. S7). The assays were applied to bulk RNA samples collected from select locations in two different ocean basins (North Atlantic, NA; South China Sea, SCS; Fig. 4A; Supp. Table S3) in which field-based microscopy observations of symbiotic *Richelia* RintRC01 were reported previously [44]. Additionally, we mined a publicly available meta-transcriptome from the subtropical North Pacific (NP) gyre (Fig. 4B) for identifying the diel expression patterns of RintRC_3892, RintRC_4851, and RintRC3409 [37]. In the meta-transcriptome dataset, all three SulP-like proteins were expressed, and RintRC_3892 clearly showed a diel periodicity of highest expression shortly after sunrise (Fig. 4B). Likewise, we consistently detected gene transcripts of RintRC_3892 and RintRC_4851 in both the NA and SCS, and RintRC_3892 was usually higher in normalized gene expression (Fig. 4C).

**Fig. 4.**
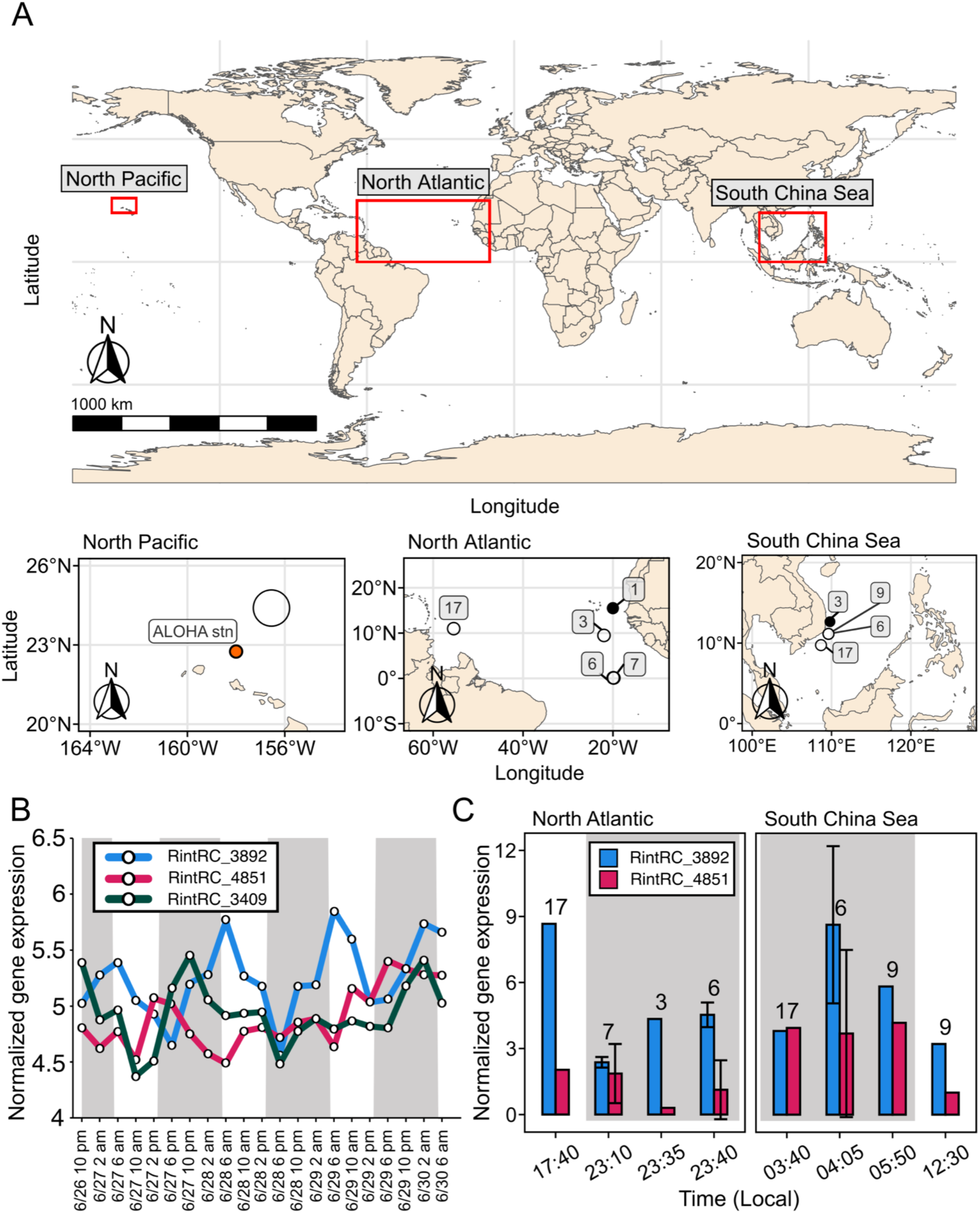
Expression of genes encoding SulP-like proteins of *Richelia* RintRC01 in wild populations. **(A)** Overview of the global map (top) and locations (below) from which samples for estimating expression of RintRC_3892 and RintRC_4851 by RT-qPCR were collected (North Atlantic, NA; South China Sea, SCS). Transcripts for RintRC_3892, RintRC_4851, and RintRC_3904 were taken from a publicly available transcriptome in the North Pacific (NP), [37]. Filled circles indicate stations in NA and SCS where neither SulP-like encoding genes were detected. **(B)** Diel patterns for normalized gene expression of RintRC_3892, RintRC_4851 and RintRC_3904 from the meta-transcriptome study in the NP [37]. **(C)** Results of normalized gene expression for RintRC_3892 and RintRC_4851 from stations in the NA and SCS collected at different times; the *secB* gene was used for normalization. The numbers above the bars indicate the station number. For stations which had multiple samples from different depths, the normalized gene expression was averaged (see Suppl. Table S3). White and gray boxes in B and C indicate the light and dark periods, respectively.

## DISCUSSION

Oceans are an important reservoir of DIC [4] that is used by marine phytoplankton to perform photosynthesis. In the biological C pump, marine diatoms, especially DDAs, make an important contribution by forming large and expansive blooms which reduce a significant amount of inorganic carbon and subsequently rapidly sink as particulate carbon [18, 19, 27, 45, 46, 47]. The DDAs also fix high amounts of N_2_, and release some of the fixed N to the surround, thus supporting primary production. In fact, under bloom conditions, the amount of new N released by DDAs to the surround can exceed the nitrate flux from below the photic zone [26], and thus how the DDAs sustain such high rates of N_2_ fixation is an important and open question. In heterocyst-forming cyanobacteria, energy derived from photosynthesis in the vegetative cells supports the N_2_ fixation in the heterocysts, hence the concentration and transport of inorganic carbon is essential to consider.

The *Richelia* spp. genomes possess the necessary genes for photosynthesis, and therefore both partners of DDAs are photosynthetic. Moreover, high rates of carbon fixation have been reported when DDAs are present [19, 26, 48]. To investigate the capabilities of inorganic carbon uptake by the endosymbiotic *Richelia* spp. living in different cellular locations in their respective diatom hosts, one must consider the CO_2_-concentration mechanism (CCM) of their respective diatom hosts. However, the CCM of the DDAs is unknown, and thus we considered that they perhaps have a CCM like model diatoms. For example, most diatoms take up CO_2_ from the external environment by passive diffusion because its plasma membrane is highly permeable to CO_2_ [49]. In addition, diatoms have evolved a CCM that enables them the ability to take up bicarbonate present in seawater by active membrane transporters of the solute carrier (SLC) 4 type transporters, which are Na^+^-dependent transporters that function as plasma as well as chloroplast membrane bicarbonate transporters [50]. Diatom hosts of the DDAs might contain, as in the rest of marine diatoms, a highly efficient CCM taking up CO_2_ and bicarbonate into the cytoplasm. Under such a scenario, the CCM of the host diatom could generate a dynamic concentration range of both CO_2_ and bicarbonate across its cell as predicted in model diatoms. For example, the concentration of CO_2_ is similar in both the cytoplasm and the periplasm of the diatom (9 and 10 μM, respectively), but the concentration of bicarbonate is an order of magnitude lower in the cytoplasm (0.4 mM) compared to the periplasm (2 mM) [11] (Fig. 5). Hence, this could impact the DIC acquisition strategies of the endobionts which reside in the two different locations: cytoplasm vs. periplasm. In fact, such a scenario might potentially create a DIC competition between the diatom and cyanobacterial partners. This study focused on the identification of transporters which could mediate the uptake of bicarbonate by the diatom endosymbionts that reside in the cytoplasm and the periplasm, i.e., the SulP-family transporters of *Richelia euintracellularis* HH01 and *Richelia intracellularis* RC01, respectively.

**Fig. 5.**
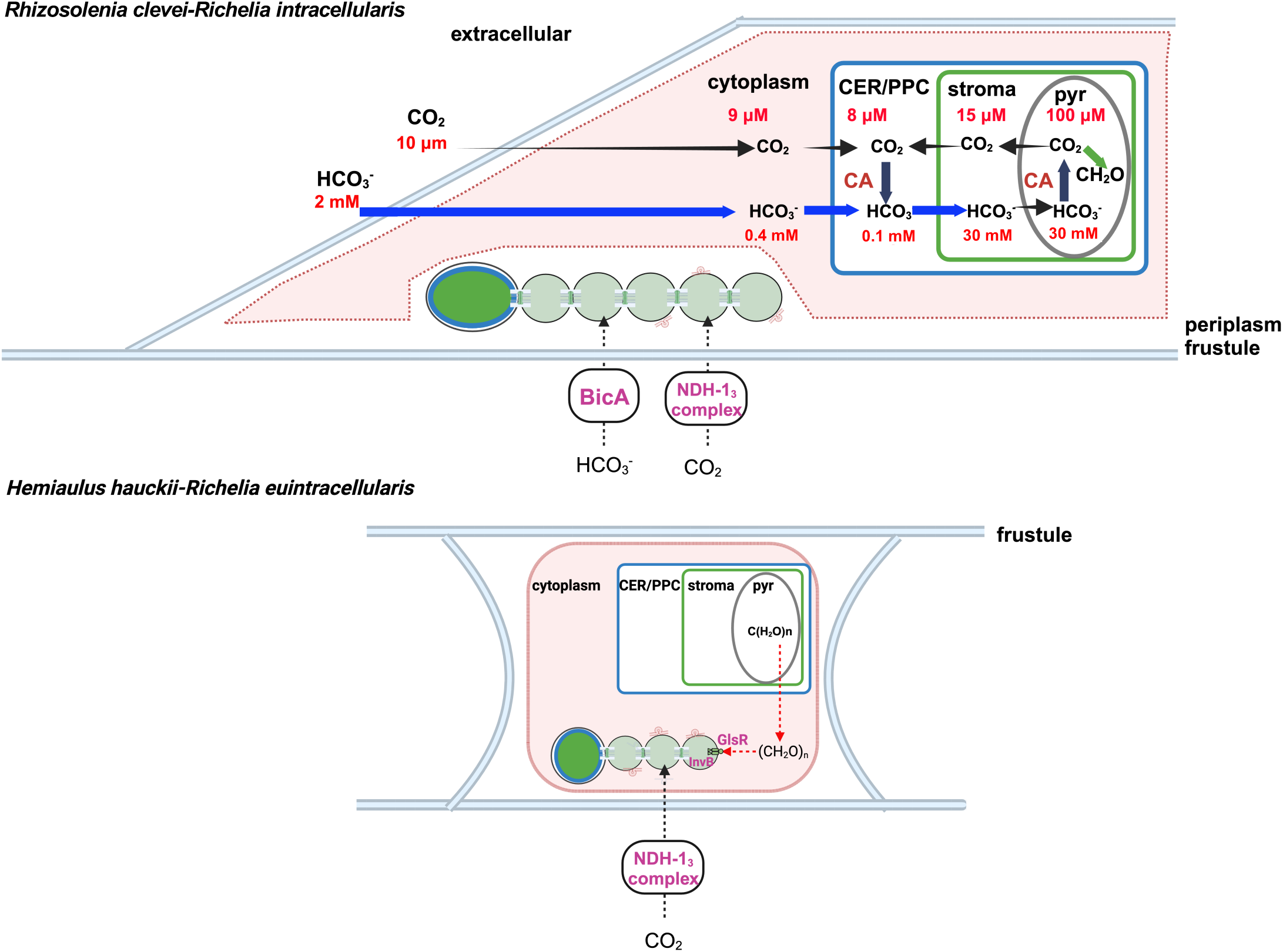
Overview of the inorganic carbon acquisition in two common Diatom Diazotroph Associations (DDAs). *Richelia intracellularis* (RintRC01) resides in the periplasm of *Rhizosolenia clevei* diatoms (top), and *Richelia euintracellularis* (ReuHH01) lives as a true endobiont in the cytoplasm of *Hemiaulus hauckii* diatoms (bottom) [20, 21]. The concentrations of inorganic carbon (CO_2_, HCO_3_^-^) vary tremendously within a diatom and results from the passive diffusion (black arrows) and active transport (blue arrows) via the hypothesized ‘chloroplast pump’ mechanism. Predicted concentrations of CO_2_ and HCO_3_^-^ shown as described in [11] only in *R. clevei* for simplicity; the same concentrations are presumed here for *H. hauckii.* Carbonic anhydrases (CA) also function in conversion of inorganic carbon; the CCM in DDAs is unknown, and assumed to function similar to that shown which is based on model diatoms [49]. As presented in this work, a SulP family BicA transporter was identified for RintRC01, with an expected *K*_0.5_ (about 626 µM) for bicarbonate (periplasm concentration similar to that in the extracellular medium, about 2 mM), while no evidence for a BicA transporter for the true endobiont ReuHH01 was obtained. Combined our results suggest that the cellular location of the *Richelia* endobionts influences both the function and affinity for inorganic carbon transport. Also shown for *H. hauckii-R. euintracellularis* is the recent demonstration for the role of organic carbon (i.e., sugars) supplied by the host diatoms via a GlsR transporter and a neutral invertase (InvB) [52]. Previous work has shown the presence of membrane vesicles formed on the cell envelope of both *Richelia* strains which could potentially function in metabolite transfer between partners [20, 21]. The following abbreviations apply: pyr, pyrenoid; CER/PPC, chloroplast endoplasmic reticulum/periplastidial compartment. Figure has been made in Biorender.

We initially found that SulP-like proteins vary in number and completeness in *Richelia* spp. genomes. Moreover, the results were also consistent in the 15 environmental MAGs reported to date. Our results have unequivocally shown that the genome of *R. intracellularis* RC01 [24], which is the periplasmic symbiont of *Rh. clevei*, bears a gene (RintRC_3892) encoding a SulP-type bicarbonate transporter. The affinity of this transporter for bicarbonate (*K*_0.5_ about 626 µM) is relatively low compared to that of high affinity bicarbonate transporters of cyanobacteria, such as the ABC transporter Cmp (K*_D_* for bicarbonate of the CmpA protein, 5 µM) [51] or the Na^+^- dependent monocomponent SbtA transporter (K*_m_* for bicarbonate about 2 µM in *Synechococcus* sp. PCC 7002) [30]. However, the affinity of RintRC_3892 for bicarbonate is in the range of the closely related *Synechcoccus* sp. PCC 7002 BicA, which is a known low-affinity but high-flux transporter (K*_m_*, 217 µM) [30]. Importantly, the affinity of RintRC_3892 is within the concentration range of bicarbonate expected in the periplasm of diatoms (e.g. 2 mM) [11]. The sodium dependency of RintRC_3892 is also similar to that of *Synechococcus* sp. PCC 7002 BicA (compare Fig. 3B to data in [30]). These observations are consistent with the close phylogenetic position of RintRC_3892 and BicA from different cyanobacteria, suggesting that RintRC_3892 from the periplasmic endobiont is a genuine BicA protein. Moreover, we consistently detected the expression of RintRC_3892 in environmental samples from two ocean basins (NA and SCS), and in the case of the North Pacific transcriptome study [37], its expression pattern follows a diel periodicity which is expected for an actively photosynthesizing organism.

The *Richelia* endobionts (periplasmic *R. intracellularis* RC01 and cytoplasmic *R. euintracellularis* HH01) appear to lack any high-affinity bicarbonate transporter, Cmp or SbtA [25]. Our finding that the genome of *R. intracellularis* RC01 encodes a BicA protein, which shows relatively low affinity for bicarbonate, is therefore of interest, since this transporter is well suited to take up concentrations of bicarbonate (about 2 mM) in the marine environment. In contrast, the possible lack of any bicarbonate transport protein in *R. euintracellularis* ReuHH01 may reflect a relatively low availability of bicarbonate in the cytoplasm of the diatom. These latter results explain, in turn, the important role of the DDA host (e.g., *Hemiaulus*) supporting the physiology of the cytoplasmic symbiont (ReuHH01) with organic carbon (19, 52).

Our results failing to show complementation of the ι15 mutant or bicarbonate uptake activity by the other tested *Richelia* SulP-like proteins, two from RintRC01 and two from ReuHH01, including a fragmented protein, do not provide evidence that these proteins function in bicarbonate transport. Moreover, in our phylogenetic analysis, none of these proteins are as close to the BicA proteins as RintRC_3892 (Fig. 1). However, we cannot discount that these proteins function under different conditions than those tested in this work or have a different function altogether. Evidence of the importance of RintRC_4851 is found in our field expression analyses as it was consistently detected in all three environmental datasets (NP, NA and SCS). Additionally, both RintRC_4851 and RintRC_3409 were identified in the field transcriptome study in the North Pacific with an obvious dynamic pattern of higher expression of RintRC_3409 in the middle of the dark periods, and RintRC_4851 showing higher expression during the late photoperiods. A study on another important diazotrophic cyanobacteria, *Trichodesmium*, demonstrated that uptake, affinities, and half-saturations (K_0.5_) for bicarbonate can vary diurnally [53], which may imply expression of different transporters.

Unlike the two endosymbionts studied here, the external facultative symbiont *R. rhizosoleniae* CalSC01 contains a high affinity bicarbonate transporter (SbtA), which is not present in the endosymbionts [25]. It can be suggested that this extra transporter is involved in bicarbonate uptake to circumvent the possible competitive situation of living in the phycosphere of its diatom host or when living unattached and free. Thus, our results are consistent with the hypothesis that these cyanobacterial endosymbionts differ in their carbon metabolic dependency with their respective hosts as a result of their adaption to an endosymbiotic life, especially in the case of the cytoplasmic endosymbiont *R. euintracellularis* (Fig. 5).

## ACKNOWLEDGEMENTS

We acknowledge support from The Swedish Research Council (Vetenskapsrådet), grant no. 2018-04161 to RAF and EF. Additional funding was provided by research grant from Knut and Alice Wallenberg Foundation to RAF. RAF thanks David Hutchins and Joseph Montoya for their invaluable support for field sampling and the captains and crew of the R/V Atlantis and R/V Falkor for providing opportunities for sample collections in the North Atlantic (2018) and South China Sea (2016), respectively; additionally, RAF acknowledges Andreas Novotny, Philip Ley, Massimo Pernice, and Andrea Caputo for field work; and Philip Ley, Klara Freed, Hanna Wolff, and Linnéa Ström for help with sample processing. MNM acknowledges María de Gracia Benítez-Eslava for help with the cyanobacterial cultures.

## AUTHOR CONTRIBUTIONS

MNM, EF and RAF conceived the study, designed experiments and drafted the manuscript. MNM, RRG and EF performed lab experiments. SB and RAF performed expression analysis in environmental samples. LLM and MH provided strains and genetic tools. All authors read, commented and approved the manuscript.

## ADDITIONAL INFORMATION

**Supplementary information.** The online version will contain supplementary material.

**Correspondence** and requests for materials should be addressed to Mercedes Nieves-Morión.

