## Supplementary figures, tables and methods for "Using gene complementation to identify a SulP-family bicarbonate transporter in an N2-fixing cyanobacterial endosymbiont of an open ocean diatom"

Content of each file:

Figure S1. SulP-like proteins from *Richelia* spp.
Figure S2. Cloning strategy to express *Richelia* genes in the *Synechocystis* Δ5 mutant.
Figure S3. DNA sequence covering the RintHH_3960-3970-3980-3990 *sulP*-like gene fragments.
Figure S4. PCR analysis showing the presence of the *Richelia* constructs in the *Synechocystis* Δ5 mutant transformants.
Figure S5. Growth of WT *Synechocystis* and the transformed Δ5 mutant in air levels of $CO_2$ supplemented or not with bicarbonate.
Figure S6. Uptake of $^{14}C$-bicarbonate at pH 7 and pH 6 by WT *Synechocystis* and the Δ5 mutant transformed with the indicated constructs.
Figure S7. Summary of the specificity testing of the oligonucleotides used in the RT-qPCR assays for estimating the expression of the SulP-like transporters in field samples by BLASTn analyses (details of databases in Supp. Methods).
Table S1. List of the oligonucleotides used in the construction of plasmids containing the symbiotic *Richelia* genes.
Table S2. Summary of oligonucletodes used in the RT-qPCR assays.
Table S3. Summary of results from the RT-qPCR assays to estimate the expression of RintRC_3892 and RintRC_4851.

### Figure S1

**Fig. S1. SulP-like proteins from *Richelia* spp.** (*Top*) BlastP analysis was performed using as query the SulP-type bicarbonate transporter BicA of *Synechococcus* sp. PCC 7002 or, when indicated, the SulP-like protein Alr1633 from *Anabaena* sp. PCC 7120. BlastP expect values are shown in parenthesis. (*Bottom*) Scheme showing the RintHH_3960-3970-3980-3990 genomic region.

| **Homologues in *R. rhizosoleniae* SC01** | **Homologues in *R. intracellularis* RC01** | **Homologues in *R. euintracellularis* HH01** |
|---|---|---|
| Ga0265390_10577 (0.0)<br><br>Ga0265390_11353 (0.0)<br><br>Ga0265390_12504 ($1e^{-18}$)<br><br>Ga0265390_13451 ($6e^{-44}$) (0.0, compared to Alr1633) | RintRC_3892 (0.0)<br><br>RintRC_4851 (0.0)<br><br>RintRC_3409 (0.0, compared to Alr1633) | RintHH_3960 ($4e^{-12}$)<br>RintHH_3970 ($e^{-103}$)<br>RintHH_3980 ($3e^{-21}$)<br>RintHH_3990 ($5e^{-32}$)<br><br>RintHH_20770 (5e-43) (0.0, compared to Alr1633) |

Gene possibly split into four fragments

Genomic neighborhood of RintHH_3970 (red color)

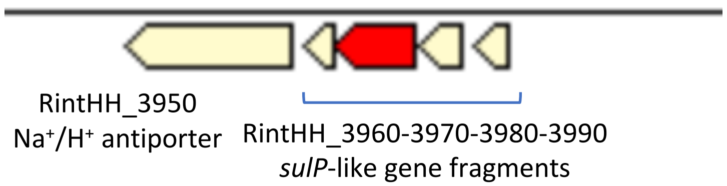

### Figure S2

**Fig. S2. Cloning strategy to express *Richelia* genes in the *Synechocystis* △5 mutant.** The restriction sites used for cloning were: NdeI and SphI for RintRC_3892, NdeI and XhoI for RintRC_4851, BamHI and BamHI for RintRC_3409, RintHH_3990-60 and RintHH_20770. The RintRC_3892 gene was PCR-amplified and cloned in pSpark prior to transfer to the NdeI/SphI site of pNRSD_PcpcB_Ery.

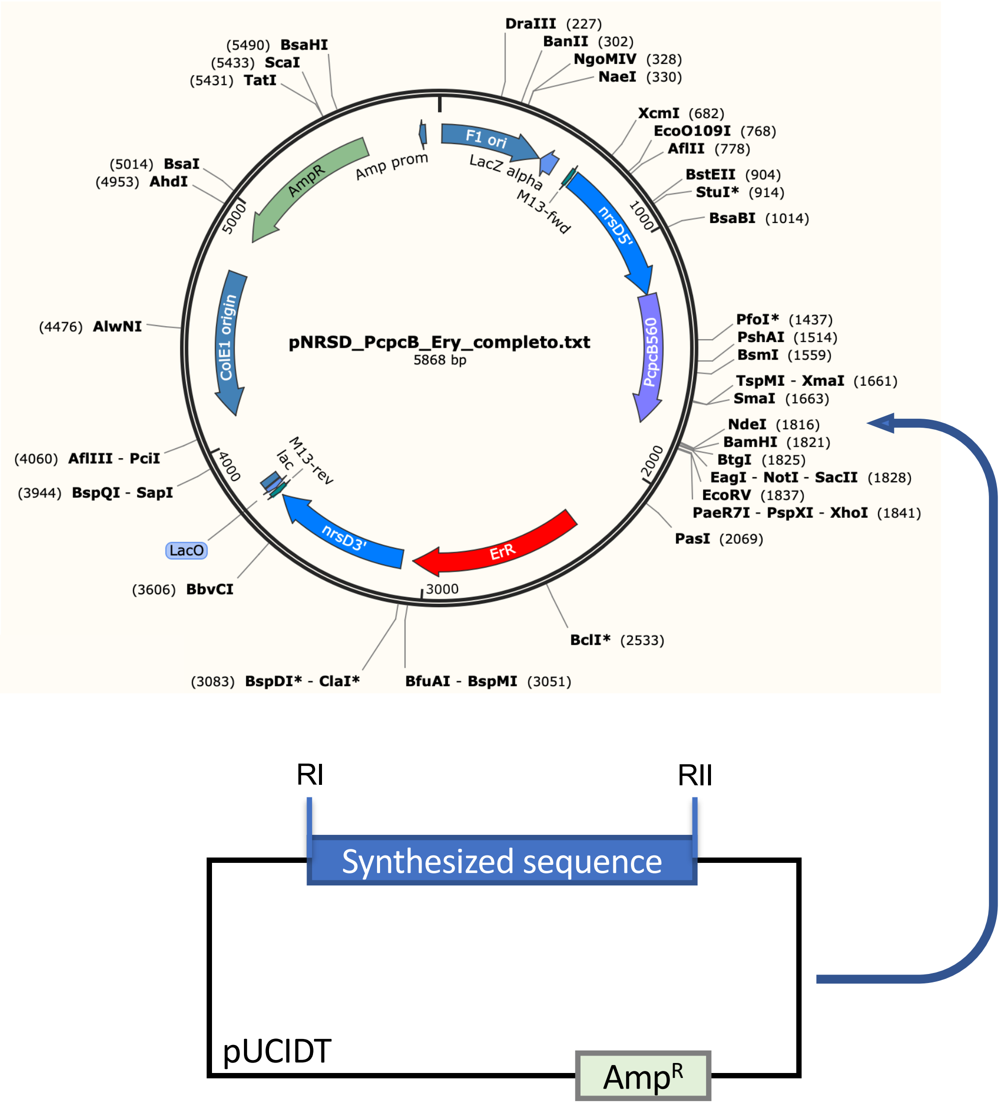

### Figure S3

**Fig. S3. DNA sequence covering the RintHH_3960-3970-3980-3990 *sulP*-like gene fragments.** The RintHH_3960-3970-3980-3990 genes (green, blue, red and magenta letters, respectively) were cloned together with their natural intervening DNA sequences (*upper panel*). The transcript from the cloned DNA fragment should be translated producing four polypeptides that align with consecutive fragments of BicA from *Synechococcus* sp. PCC 7002 (*lower panel*).

```
GGATCCATGCAAATTTTGAATCGCATCCATTTTAGAAATCTTCGTGGTGACATCTTTGGAGGTTTAACCTCAGCGATTA
TCTCCTTGCTCTTAGCTATTGCATTCAGTGTTGCTTCTGGGATGAGACCAATATCAGGTGTATATGGTGCTGTGATAAT
AGGTTTATTTGCAGCATTATTTGGTGTTACACCAACCCTAATAATTTCCGAACCCACTGGGCCAATGACTGTCATCATG
ACTGGTGTTATAGCTTCTATGATAGCAAAGGATACAGAGTAGCGTGGCAATGGCATTCACGGTGGTAATCTTAGCAGGT
ATGATTTAATTACTGTTTGGTATATTTAAACCAGGCAAATATATTACCTTAATGCCTTATAGTGTTATTTCTGGCTTCA
TGTCGGGAATTGTTTCAATTCTAGTGATTTTACAAATTGCCCCATTGGTGGGCGAACAAGTACCAAAAGGTGGGGTAAT
TGGCACTATTATTAGCAATATTCCAACCTTAATTACTAATATTAATGCCCCTGATTTGGTATTAGGAGGGTTAACAATC
ACTATTTTATTCCTAACACCATCAAAATTGAAGTATTTCTTTCCTCCACATTTAATTGCATTAATTATTGGTACTCTAG
TTTATATAACTGTTTTGCAACATCCTGAGATAGCAAGAATACCAGAAATCCCTGCAGAATGGCCAAAACTACAATTACC
GTATTTCACACCCGGGCAAATAACTTGCTATTAGATAGTATTGTTTTGGCAATAATGGGGTGTATTGATACTCTTTTAA
CCTCTGTAATTGCAGATAGTTTAACCCGTGTTGAGCATAAATCTAATAAAGAACTAATTTGTCAGGACATTGCTAACTT
AATTTCTGGCTTGTTTGGGGGTCTACCTGGTGCAGATGCAACTATAGGAACTATAGTTAACATTCAAACAGGAGCTAAA
ACTGCACTATCAGGAGTAACTCGTAGTTTAGTATTAATAATAGTAATTTTGAGTGCTGCAAGACTAGCTCAAAATATTC
CCATGTCTGTATTGACTGGTATTGCCTTAAAAGTGGGTCTTGATATCCTAGATTGGAATTTCCTCAAGTGTGCTCACAA
AGTATCACTTAAGGGTGCGCTAATTATGTATGATGTATTATTTTTAACAATATTTGTAGATTTAATTGTTGCTGTAGGT
GTAGGTTTATTCATAGCAAATATTTTGACTATTGAGCATCTTTTTAACCTTCAATCAAAAGAAGTTAAAACTATTAGTG
ATACTGATGAAAAAATTAATCTAACTAATATAGAGAGGTCTTTACTTGAACAAGTAAATGGAAGAATACTGTTATTTTA
CCTTAATAGTCCTATGATATTTAGGGTGGCAAAAGCAATTTCTCGAGAGCATTCAGCGATGAGAGATGCAGATGCTCTC
ATAATAGATTTAAGTGATGTACCGTACCAATGTTAGGAGTAACTGCTTGTTTGGCAATTGAAAACACAGTTAAAGATGG
AGTTCATAGAGGTCTGCAAGTATTTATTGTGAGTGCTGCAGGTAAGGTTAAACAGCGTTTAGAACGATTTGACTTATCA
CAAATTTTACCTCCCAACCACTTGTTGGCAAATCGTACAGAGGCACTTCAGCAAGCACTTTTTTTTGTAATAAAATACT
CAGGCTACACAAGTAGTAACATTCTAGCATTGGATGATTCAACCATTTTGTAGGGATCC
```

```
Synechococcus      MQITNKIHFRNIRGDIFGGLTAAVIALPMALAFGVASGAGAEAGLWGAVLVGFFAALFGG 60
RintHH3990-60      MQILNRIHFRNLRGDIFGGLTSAIISLLLAIAFSVASGMRPISGVYGAVIIGLFAALFGV 60
                   *** *:*****:*********:*:*:* :*:**.****     :*::***::*:******

Synechococcus      TPTL-ISEPTGPMTVVMTAVIAHFTASAATPEEGLAIAFTVVMMAGVFQIIFGSLKLGKY 119
RintHH3990-60      TPTLIISEPTGPMTVIMTGVIASMIAK--------------------------------- 87
                   **** **********:**.*** : *.

Synechococcus      VTMMPYTVISGFMSGIGIILVILQLAPFLGQASPGGGVIGTL-QNLPTLLSNIQPGETAL 178
RintHH3990-60      ---------DTEMSGIVSILVILQIAPLVGEQVPKGGVIGTIISNIPTLITNINAPDLVL 138
                             .  ****  ******:**::*:  * ******: .*:***::**:  : .*

Synechococcus      ALGTVAIIWFMPEKFKKVIPPQLVALVLGTVIAFFVFPPEVSDLRRIGEIRAGFPELVRP 238
RintHH3990-60      GGLTITILFLTPSKLKYFFPPHLIALIIGTLVYITVL--QHPEIARIPEIPAEWPKLQLP 196
                   .  *::*::: *.*:* .:**:*:**::**:: : *:  :  :: ** ** * :*:*  *

Synechococcus      SFSPVEFQRMILDAAVLGMLGCIDALLTSVVADSLTRTEHNSNKELIGQGLGNLFSGLFG 298
RintHH3990-60      YFTPGQI---------TCYMGCIDTLLTSVIADSLTRVEHKSNKELICQDIANLISGLFG 247
                    *:* ::           :****:*****:******.**:****** *.:.**:*****

Synechococcus      GIAGAGATMGTVVNIQSGGRTALSGLVRAFVLLVVILGAASLTATIPLAVLAGIAFKVGV 358
RintHH3990-60      GLPGADATIGTIVNIQTGAKTALSGVTRSLVLIIVILSAARLAQNIPMSVLTGIALKVGL 307
                   *: **.**:**:****:*.:*****:.*::**::***.** *: .**::**:***:***:

Synechococcus      DIIDWSFLKRAHEISPKGALIMYGVILLTVLVDLIVAVGVGVFVANVLTIERMSNLQSEK 418
RintHH3990-60      DILDWNFLKCAHKVSLKGALIMYDVLFLTIFVDLIVAVGVGLFIANILTIEHLFNLQSKE 367
                   **:**.*** **::* *******.*::**::**********:*:**:****:: ****::

Synechococcus      VQTVSDADDNIRLTTTEKRWLDEGQGRVLLFQLSGPMIFGVAKAIAREHNAMGDCDALVF 478
RintHH3990-60      VKTISDTDEKINLTNIERSLLEQVNGRILLFYLNSPMIFRVAKAISREHSAMRDADALII 427
                   *:*:**:*::*.**. *:  *:: :**:*** *..**** *****:***.** *.***::

Synechococcus      DIGEVPH---MGVTASLALENAIEEALDKERQVYIVGAAGQTRRRLEKLKLFKRVPPDKC 535
RintHH3990-60      DLSDVPYQCMLGVTACLAIENTVKDGVHRGLQVFIVSAAGKVKQRLERFDLSQILPPNHL 487
                   *:.:**:   :****.**:**::::.:.:  **:**.***:.::***::.* : :**::

Synechococcus      LMSREEALKNAVLGIYPHLADGVTAPSSEMG-------      566
RintHH3990-60      LANRTEALQQALFFVIKYS--GYTS-SNILALDDSTIL      522
                   * .* ***::*:: :  :   * *: *. :.
```

### Figure S4

**Fig. S4. PCR analysis showing the presence of the *Richelia* constructs in the *Synechocystis* Δ5 mutant transformants.** PCR analysis was performed with the primers indicated in **(A)** and DNA was isolated from the *Synechocystis* strain indicated in **(B)**: WT or the Δ5 mutant transformed with the indicated gene and grown as described in Materials and Methods. X symbols indicate approximate sites of recombination with the *Synechocystis* chromosome. RI and RII, restriction endonuclease cutting sites (see Fig. S2). L, 1-kb DNA ladder (Biotools) above the 1-7 next to the gel.

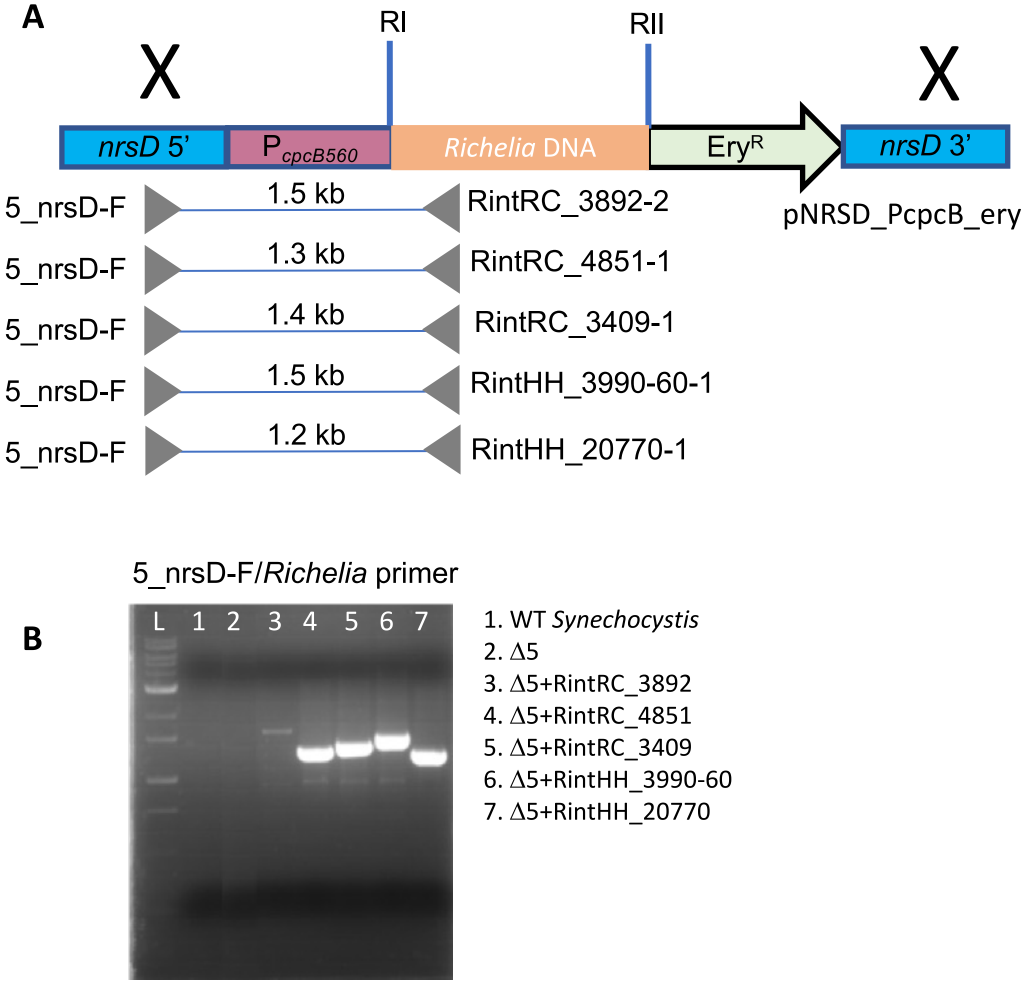

### Figure S5

**Fig. S5. Growth of WT *Synechocystis* and the transformed Δ5 mutant in air levels of $CO_2$ supplemented or not with bicarbonate.** Photoautotrophic growth at 30 ºC under the indicated conditions: **(A)** bubbled cultures, (**B)** shaken cultures. 1, *Synechocystis* WT; 2, Δ5; 3, +RintHH_3990-60; 4, +RintHH_20770; 5, +RintRC_3409; 6, +RintRC_4851; 7, +RintRC_3892.

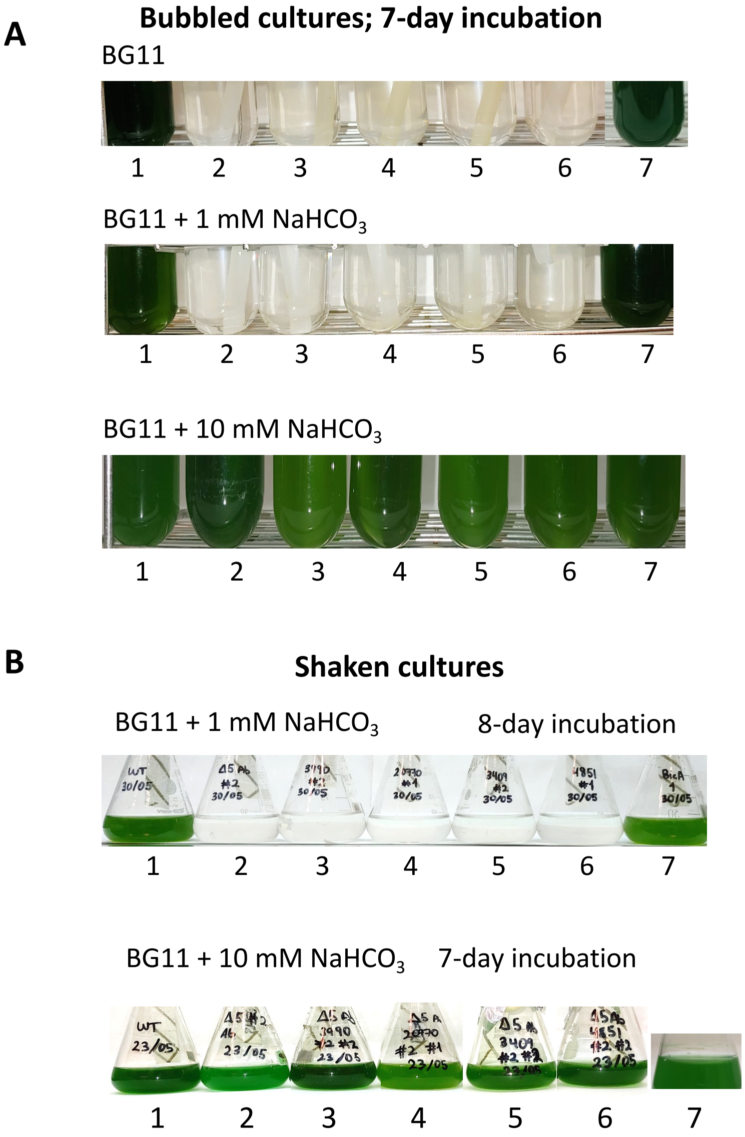

### Figure S6

**Fig. S6. Uptake of $^{14}C$-bicarbonate at pH 7 and pH 6 by WT *Synechocystis* and the Δ5 mutant transformed with the indicated constructs.** Uptake was tested with 1 mM $NaH^{14}CO_3$ in BG11/2 medium with 25 mM TES-KOH (pH 7) or MES-KOH (pH 6) buffer; final $Na^+$ concentration, about 18.5 mM. Assays were 1-min in duration.

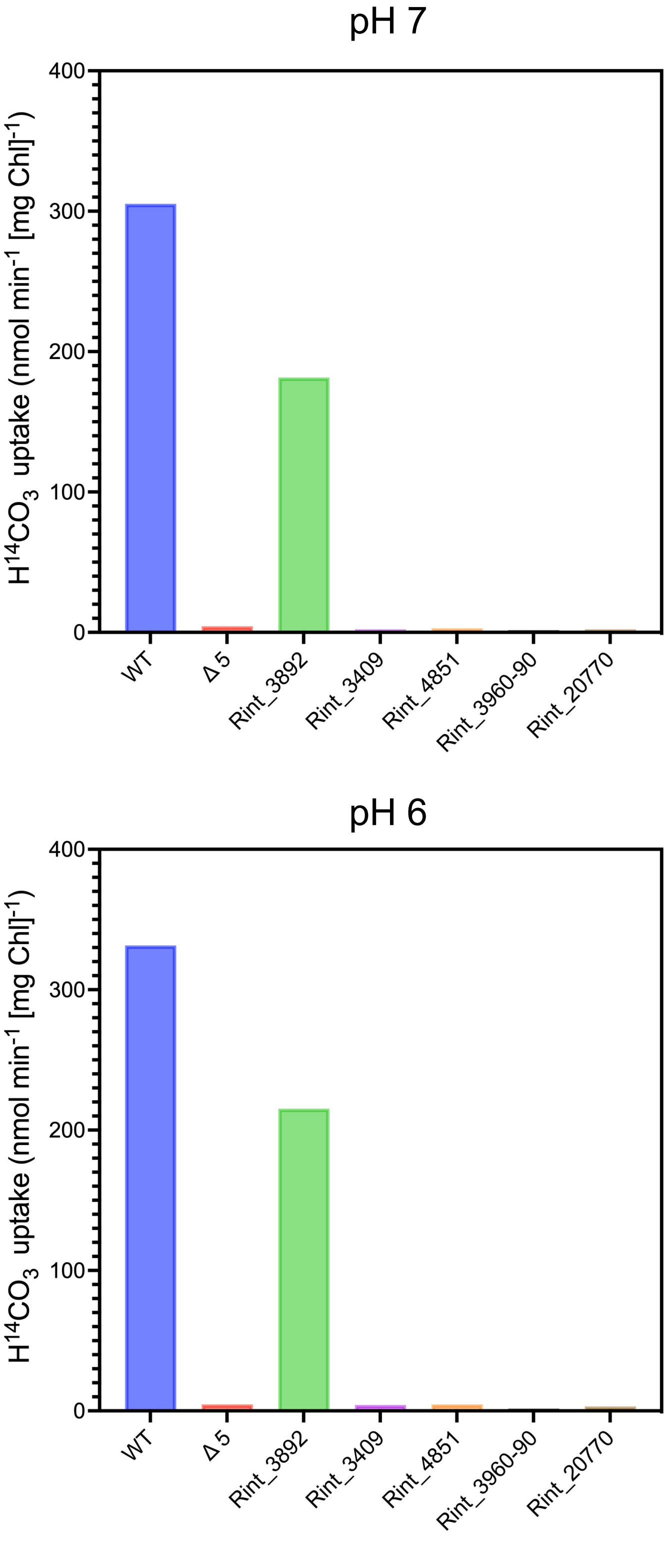

### Figure S7A

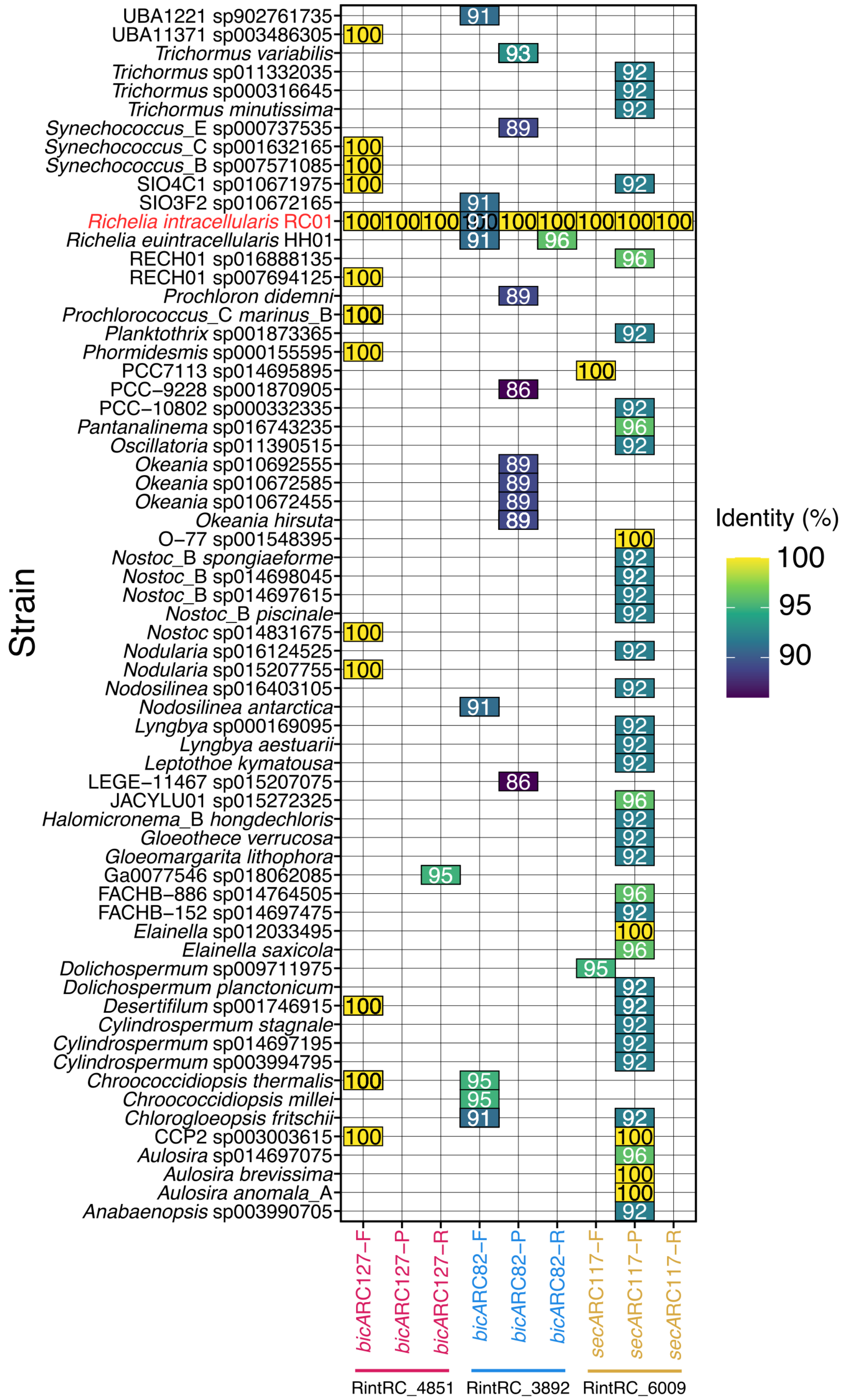


**Fig. S7**. **Summary of the specificity testing of the oligonucleotides used in the RT-qPCR assays for estimating the expression of the SulP-like transporters in field samples by BLASTn analyses (details of databases in Supp. Methods).** Two oligonucleotide sets were designed based on RintRC_3892 and RintRC_4891; an additional oligonucleotide set was developed for normalizing expression based on *secA* (RintRC_6009) (Suppl. Table S2). **(A)** Results of BlastN analyses using a local database containing genomes of cyanobacteria originating from RefSeq and GenBank (9,680,513 sequences) based on phylogenetic classification according to the Genome Taxonomy Database (GTDB version R207). Only alignments that used the whole query sequence were used in the analyses. The x-axis indicates the primer used as a query (bicARC127, RintRC_4851; bicARC87, RintRC_3892; secARC117, RintRC_6009). The primer names are in the structure of 'gene name', 'length of amplified sequence' and forward (F), reverse (R) or probe (P). **(B)** Results of BlastN analyses from Fig S7A were filtered for *Richelia* strains, which also included environmental *Richelia* MAGs. **(C)** BlastN alignments for oligonucleotides designed to detect RintRC01_3892 as queries against RintRC_4851 (top) and RINTHH_3990 (middle, *bic*ARC82-F; bottom, *bic*A82-R). Sequence similarities are shown in Fig S7B.

### Figure S7B

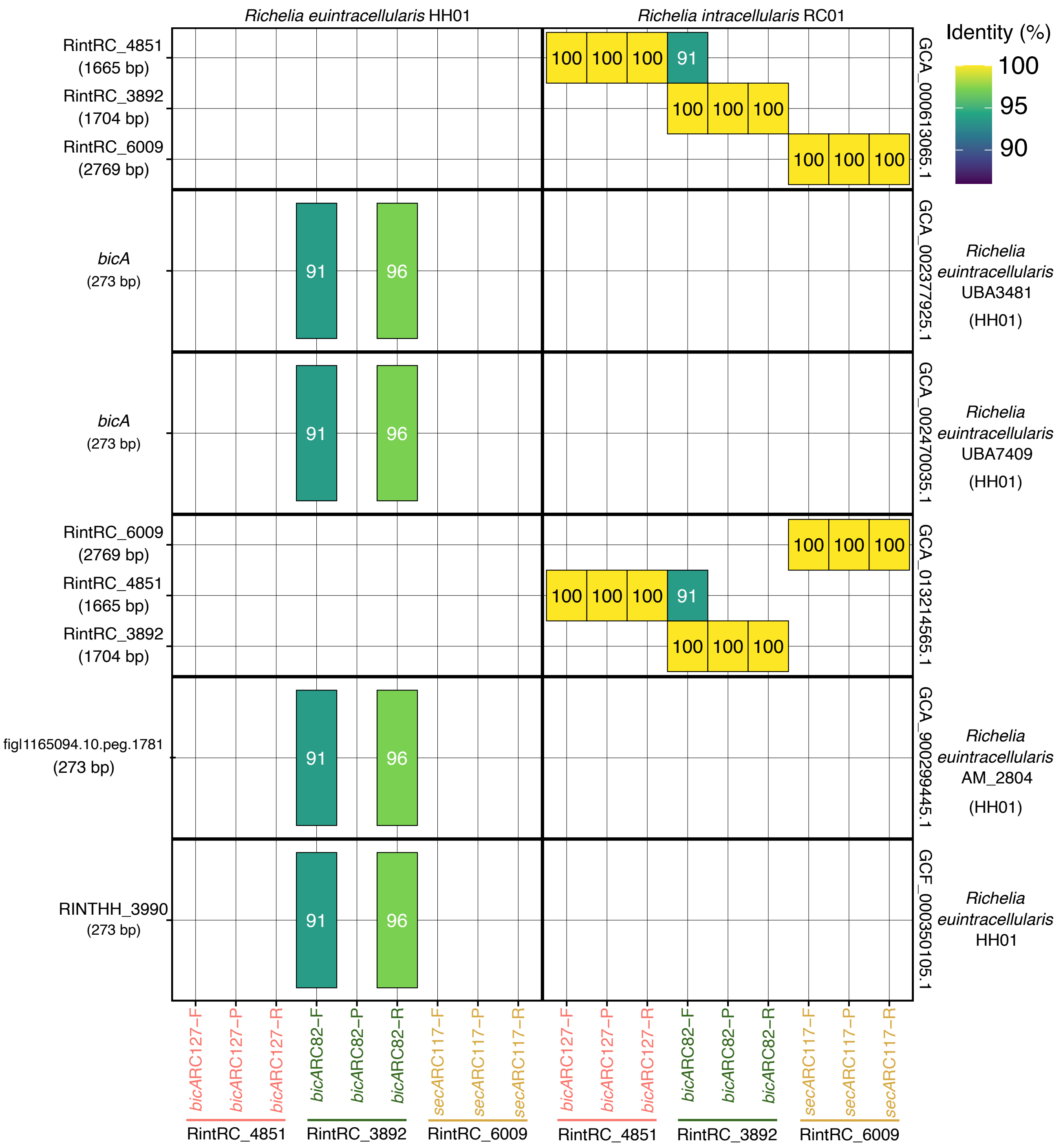

Richelia euintracellularis HH01
Richelia intracellularis RC01
Identity (%)
100
95
90
RintRC_4851 (1665 bp)
RintRC_3892 (1704 bp)
RintRC_6009 (2769 bp)
GCA_000613065.1
bicA (273 bp)
GCA_002377925.1
Richelia euintracellularis UBA3481 (HH01)
bicA (273 bp)
GCA_002470035.1
Richelia euintracellularis UBA7409 (HH01)
RintRC_6009 (2769 bp)
RintRC_4851 (1665 bp)
RintRC_3892 (1704 bp)
GCA_013214565.1
fig|1165094.10.peg.1781 (273 bp)
GCA_900299445.1
Richelia euintracellularis AM_2804 (HH01)
RINTHH_3990 (273 bp)
GCF_000350105.1
Richelia euintracellularis HH01
100 100 100 91
100 100 100
100 100 100
91 96
91 96
100 100 100
100 100 100 91
100 100 100
91 96
91 96
bicARC127–F
bicARC127–P
bicARC127–R
bicARC82–F
bicARC82–P
bicARC82–R
secARC117–F
secARC117–P
secARC117–R
RintRC_4851
RintRC_3892
RintRC_6009

### Figure S7C

```
>JHFCPCOE_05849 Bicarbonate transporter BicA [GCA_000613065.1
d__Bacteria;p__Cyanobacteria;c__Cyanobacteriia;o__Cyanobacteriales;f__Nostocaceae;g__Richelia;s__Richelia
intracellularis_B]
Length=1665

 Score = 31.9 bits (34),  Expect = 6.0
 Identities = 20/22 (91%), Gaps = 0/22 (0%)
 Strand=Plus/Plus

Query  1    GGTTTATTTGCAGCGTTGTTTG  22
            |||||||||||||| || ||||
Sbjct  169  GGTTTATTTGCAGCTTTATTTG  190
>IPCPKDNE_01849 Bicarbonate transporter BicA [GCF_000350105.1
d__Bacteria;p__Cyanobacteria;c__Cyanobacteriia;o__Cyanobacteriales;f__Nostocaceae;g__Richelia;s__Richelia
intracellularis]
Length=273

 Score = 31.9 bits (34),  Expect = 6.0
 Identities = 20/22 (91%), Gaps = 0/22 (0%)
 Strand=Plus/Plus

Query  1    GGTTTATTTGCAGCGTTGTTTG  22
            |||||||||||||| || ||||
Sbjct  154  GGTTTATTTGCAGCATTATTTG  175
>IPCPKDNE_01849 Bicarbonate transporter BicA [GCF_000350105.1
d__Bacteria;p__Cyanobacteria;c__Cyanobacteriia;o__Cyanobacteriales;f__Nostocaceae;g__Richelia;s__Richelia
intracellularis]
Length=273

 Score = 41.9 bits (45),  Expect = 0.012
 Identities = 24/25 (96%), Gaps = 0/25 (0%)
 Strand=Plus/Minus

Query  1    CACCAGTCATGATAACAGTCATTGG  25
            ||||||||||||| |||||||||||
Sbjct  238  CACCAGTCATGATGACAGTCATTGG  214
```

**Table S1. List of the oligonucleotides used in the construction of plasmids containing the symbiotic *Richelia* genes**. Introduced restriction enzyme cutting sites are underlined.

| **Primer name** | **Sequence (5´ to 3´)** |
|---|---|
| RintRC_3892-5 (ClaI & NdeI sites) | GGCATCGATAAGGAATTATAACATATGCAACTTTTGAATCGCATCCAC |
| RintRC_3892-6 (PvuI & SphI sites) | AAACGATCGAAGCATGCTCTGACTATGAGTTTGTGGGCA |
| RintRC_3892-2 | CAGGTGAATCATCTATGGGTGC |
| RintRC_3892-7 | CCGATACCCGACATGAAGC |
| RintRC_4851-1 | ACGTGAGGCGGAAACTCCAA |
| RintRC_3409-1 | AAGGCGGTCAAAACAGCGAT |
| RintHH_3990-1 | AATGCCATTGCCACGCTACT |
| RintHH_20770-1 | CAAAATGTCGGCTCTCGCGT |
| PcpcB560_F | ATCGAATTCCACCTGTAGAGAAGAGTC |
| 5_nrsD-F (BssHI site) | AAGCGCGCCTTTCACTGCTTGCGGAACC |

**Table S2. Summary of oligonucletodes used in the RT-qPCR assays.** RintRC_3892 and RintRC_4851 were designed to amplify the SulP-like Proteins, and RintRC_6009 was designed to amplify the secA gene. Sequences were taken from RintRC_01 draft genome (Genbank Assembly: GCA_ 000613065.1)

| **Target** | **Forward 5′ to 3′** | **Probe 5′ to 3′** | **Reverse 5′ to 3′** |
|---|---|---|---|
| RintRC_3892 | GGTTTATTTGCAGCGTTGTTTG | TACACCCACCCTGATTTCCGAACCTACC | CACCAGTCATGATAACAGTCATTGG |
| RintRC_4851 | GCTTCATGTCTGGGATTG | TACCTTGGCTGAACTGGGATGTCCC | GGATTGGGAGTAGTGAGATA |
| RintRC_6009 | GAAGGTGGAGTGATGGATTA | ATTCAGCCGGAAACCCAAACCCTA | TCGGATACAGCAAGAATAGG |

**Table S3. Summary of results from the RT-qPCR assays to estimate the expression of RintRC_3892 and RintRC_4851.** Expression has been normalize to secA (RintRC_6009). Details on the sample location, time of sampling, and normalized cDNA $L^{-1}$ are provided; samples that were below detection are noted as bd, samples noted as detected but not quantifiable (dnq) indicate the samples which had 1 or 2 of the 3 replicates that amplified, samples not run are noted as nr; The ocean basin use the following abbreviations: NA, North; SA, South Atlantic; SCS, South China Sea.

| Sample ID | Station | Longitude | Longitude | Ocean Basin | Local sampling time | Depth (m) | Normalized cDNA RintRC_3892 $L^{-1}$ | Normalized cDNA RintRC_4851 $L^{-1}$ |
|---|---|---|---|---|---|---|---|---|
| 681 | 1 | 15º 30′ N | 21º 29′ W | NA | 10:14 | 5 | bd | bd |
| 682 | 1 | | | | 10:14 | 40 | bd | bd |
| 683 | 1 | | | | 10:14 | 50 | bd | bd |
| 695 | 3 | 9º 29.95′ N | 22º 0.04′ W | NA | 23:35 | 5 | 4.34 | 0.30 |
| 696 | 3 | | | | 23:35 | 20 | bd | bd |
| 697 | 3 | | | | 23:35 | 42 | bd | bd |
| 714 | 6 | 0º 0.45 ´ N | 21º 59.23 ´W | NA | 23:40 | 5 | 5.15 | 0.2 |
| 715 | 6 | | | | 23:40 | 15 | 4.36 | 0.51 |
| 716 | 6 | | | | 23:40 | 30 | dnq (2) | bd |
| 723 | 6 | | | | 11:00 | 5 | 31.35 | nr |
| 724 | 6 | | | | 10:05 | 20 | 13.15 | nr |
| 725 | 6 | | | | 10:05 | 35 | 9.08 | nr |
| 731 | 7 | 1º 59.42 ´ S | 21º 59.77 ´W | SA | 23:10 | 5 | 2.54 | 0.91 |
| 732 | 7 | | | | 23:10 | 20 | 2.20 | 2.81 |
| 733 | 7 | | | | 23:10 | 40 | bd | bd |
| 608 | 17 | 10 º59.70′ N | 55º 26.78′ W | NA | 17:40 | 5 | 8.66 | 2.03 |
| 609 | 17 | 17 | | | 17:40 | 20 | dnq (2) | dnq (1) |
| 610 | 17 | 17 | | | 17:40 | 40 | bd | bd |
| 10129 | 3 | 12º 39.04′N | 109º48.280′E | SCS | 8:40 | 0 | bd | bd |
| 10062 | 6 | 11º 8.439′N | 109º37.036′E | SCS | 4:05 | 45 | bd | bd |
| 10063 | 6 | | | | 4:05 | 30 | bd | bd |
| 10064 | 6 | | | | 4:05 | 20 | 11.15 | dnq(1) |
| 10065 | 6 | | | | 4:05 | 13 | 6.10 | 6.37 |
| 10066 | 6 | | | | 4:05 | 1 | dnq(1) | nr |
| 10165 | 9 | 11º 8.439′N | 109º 37.062′E | SCS | 12:30 | 8 | 3.21 | dnq(1) |
| 10166 | 9 | | | | 12:30 | 5 | bd | nr |
| 10171 | 10 | 9º 51.2082 ´N | 107º 0.558′E | SCS | 3:55 | 1 | 5.82 | 4.17 |
| 10203 | 17 | 9º44.5968′N | 108º 40.448′E | SCS | 3:40 | 25 (1.5L) | 3.80 | 3.94 |

#### Supplementary Methods

**Whole water collections.** Water (2-2.5l) for total RNA were collected on two different field expeditions to the North Atlantic (NA; TriCoLim expedition, 8Feb-11Mar 2018), and the South China Sea (SCS; 4June-18June 2016) from the conductivity depth temperature (CTD) rosette. Water samples were collected into bleach-rinsed (10%) bottles (2.5L) from discrete depths and filtered immediately onto 0.2 µm pore size (25 mm diameter) filter using a peristaltic pump. Volumes of water filtered in the North Atlantic were 2-2.5L, while in the SCS volumes varied from 0.5L to 3L. Filters collected in the NA were placed into sterile tubes amended with RLT buffer (Qiagen RNA easy) and stored at -80 °C until extraction. Samples collected in the SCS were initially placed in RNA later, and stored at -20 °C while in the field, then sent in a dry shipper, and stored in -80 °C until extraction. A smaller subset of samples (Suppl. Table S3) was chosen for further analyses based on field microscopy observations of higher abundances ($10^1$-$10^2$ cells/L) of the *Rhizosolenia clevei-Richelia intracellularis* symbioses.

**Design of oligonucleotides for *in situ* gene expression**. The TaqMAN oligonucleotides that target the SulP-like proteins (RintRC_3892, RintRC_4851) and *secA* (RintRC_6009) gene from *Richelia intracellularis* were designed using the PrimerQuest™ Tool (IDT, Coralville, Iowa, USA) [1] and the candidate gene sequences. The suggested oligonucleotides were initially tested for cross-reactivity using BLASTn against a database that included representative cyanobacteria genomes from the National Center for Biotechnology Information (NCBI) database Refseq [2-4] and GenBank [5] based on the Genome Taxonomy Database (GTDB; v. R07-RS207) [6] which were subsequently annotated using Prokka (v. 1.14.6) [7]. The GTDB release 7 based on RefSeq 207 (R07-RS207) includes 9 680 513 cyanobacteria genomes. Genomes for which all three oligonucleotides were significantly aligned should be considered for potential cross-reactivity, i.e. non-specificity. The BLASTn analyses showed the oligonucleotides to be highly specific for each respective target from *R. intracellularis* (See Suppl. Fig 7), and thus the cross-reaction with other related non-targets in the environmental expression assays would not be expected. The primers were subsequently synthesized by IDT.

**Phylogenetic analysis of SulP-like proteins from *Richelia* spp.** Phylogenic reconstruction was performed in Phylogeny.fr (http://www.phylogeny.fr) [8] with default values using the following ID genes: *Anabaena* sp. PCC 7120 *alr1633*, JGI gene ID 637232001; *Anabaena* sp. PCC 7120 *alr1635*, JGI gene ID 637232003; *Anabaena* sp. PCC 7120 *all1304*, JGI gene ID 637231670; *Arabidopsis* molybdate transporter, Uniprot ID Q9MAX3; *Chlamydomonas reinhardtii* molybdate transporter, Uniprot ID A6YCJ2; *Chlamydomonas reinhardtii* proton/sulfate cotransporter, Uniprot ID A8J6J0; *Clostridium* molybdate transporter, Uniprot ID A0A0G3W6Q0; *Leptospira* SulP/Carbonic anhydrase,

Uniprot ID Q8F8H7; *Mycobacterium* putative sulfate transporter, Uniprot ID P9WGF7; *Neurospora* sulfate transporter, Uniprot ID P23622; *Prosthecobacter* SulP/Carbonic-anhydrase, Uniprot ID A0A7W7YHE3; *Richelia* RintHH_20770, JGI gene ID 2580948624; *Richelia* RintRC_3409, NCBI locus CDN17084;*Richelia* RintRC_3892, NCBI locus NRB09343; *Richelia* RintRC_4851, NCBI locus NRB09526; *Richelia* RrhiSC01 Ga0265390_10577, JGI gene ID 2790719465; *Richelia* RrhiSC01 Ga0265390_11353, JGI gene ID 2790720482; *Richelia* RrhiSC01 Ga0265390_13451, JGI gene ID 2790723128; *Richelia* RrhiSC01 Ga0265390_12504, JGI gene ID 2790721855; *Synechococcus* PCC 7942 nitrate transporter, JGI gene ID 637798772; *Synechococcus* PCC 7002 BicA, JGI gene ID 641612381; *Synechococcus* WH8102 BicA, JGI gene ID 637445639; *Synechocystis* PCC 6803 BicA, JGI gene ID 2515004973, Uniprot ID Q55415; *Synechocystis* PCC 6803 high-affinity sulfate transporter, JGI gene ID 2562385466; Yeast high-affinity_sulfate_transp_SUL1, Uniprot ID P38359; *Yersinia* SulP-like transporter, Uniprot ID O07488.
